# The Novel PII-Interacting Regulator PirC (Sll0944) Identifies 3-Phosphoglycerate Mutase (PGAM) as Central Control Point of Carbon Storage Metabolism in Cyanobacteria

**DOI:** 10.1101/2020.09.11.292599

**Authors:** Tim Orthwein, Jörg Scholl, Philipp Spät, Stefan Lucius, Moritz Koch, Boris Macek, Martin Hagemann, Karl Forchhammer

## Abstract

Nitrogen limitation imposes a major transition in the life-style of non-diazotrophic cyanobacteria, which is regulated via a complex interplay of regulatory factors, involving, the nitrogen-specific transcription factor NtcA and the pervasive signal processor PII. Immediately upon nitrogen-limitation, newly fixed carbon is re-directed towards glycogen synthesis. How the metabolic switch for distributing fixed carbon to either glycogen or cellular building blocks is operated was poorly understood. Here we identify from *Synechocystis* sp. PCC 6803 a novel PII interactor, PirC, (Sll0944) that controls 3-phosphoglycerate mutase (PGAM), the enzyme that deviates newly fixed CO_2_ towards lower glycolysis. PirC acts as competitive inhibitor of PGAM and this interaction is tuned by PII/2-oxoglutarate. High oxoglutarate release PirC from PII-complex to inhibit PGAM. Accordingly, PirC deficient mutant, as compared to the wild-type, shows strongly reduced glycogen levels upon nitrogen deprivation whereas polyhydroxybutyrate granules are over-accumulated. Metabolome analysis revealed an imbalance in 3-phosphoglycerate to pyruvate levels in the PirC mutant, conforming that PirC controls the carbon flux in cyanobacteria via mutually exclusive interaction with either PII or PGAM.

## Introduction

Cellular homeostasis relies on the capacity of living systems to adjust their metabolism in response to changes in the environment. Therefore, the organisms must be able to score the metabolic state and tune it in response to environmental fluctuations. It has been proposed that cyanobacteria do not extensively rely on direct environmental sensing, but rather are primarily concerned about their internal metabolic state (1). This “introvert” lifestyle requires that they constantly and precisely monitor their intracellular milieu in order to detect imbalances caused by external perturbations. The maintenance of C/N homeostasis is one of the most fundamental aspects of cellular physiology. For photoautotrophic organisms like cyanobacteria, it is essential to tightly interconnect CO_2_ fixation and N-assimilation. To fulfil this task, cyanobacteria use a sophisticated signalling network organized by the pervasive PII signalling protein. These proteins are fundamental for most free-living prokaryotes and chloroplasts of green plants (2). The PII proteins act as multitasking signal integrators that combine information on the metabolic C/N balance through interaction with the status reporter metabolite 2-oxoglutarate (2-OG) and the energy state of the cell by competitive binding of ATP or ADP. 2-OG is ideally suited as a status reporter metabolite for C/N balance, as this tricarboxylic acid (TCA) cycle intermediate represents the precursor metabolite into which nitrogen in form of ammonia is incorporated through the assimilatory reactions catalysed by the glutamine synthetase - glutamate synthase (GS/GOGAT) cycle (3).

The interaction of PII proteins with various effector molecules, as well as how they transmit these information into conformastional states that are perceived by the targets, has been elaborated in great detail (recently reviewed (3–6)). The three intersubunit clefts of the trimeric PII proteins contain intercommunicating effector molecule binding sites: there, ADP or ATP compete for occupying these sites, whereby ATP creates a ligation sphere for the effector 2-OG through a bridging Mg^2+^ ion. This results in a synergy between ATP and 2-OG, outcompeting the ligand ADP. Depending on the effector molecules bound, the large, flexible surface-exposed T-loops, that emerge from the effector binding sites, can adopt specific conformations, allowing signal receptor proteins to read out the metabolic information through protein-protein interactions (5). A variety of key metabolic enzymes, transcription factors and transport proteins use this signalling path to tune their activity in response to the metabolic state. In many cases, the different PII conformations can directly interact with target proteins such as the N-Acetyl-L-glutamate kinase, catalysing the committed step in arginine biosynthesis (7, 8), the acetyl-coA carboxylase (ACCase), catalysing the rate-limiting step in fatty acid biosynthesis (9), the phosphoenolpyruvate carboxylase (PEPC), which catalyses an anaplerotic CO_2_-fixation (10), or the glutamine-dependent NAD^+^ synthetase (NadE) (11). In addition to the regulation of enzyme activities via PII, the trimeric protein can also influence transport activities. Recent pull-down analysis revealed that PII regulates an ensemble of nitrogen transport complexes, involving the NRT nitrate/nitrite transport system, the URT urea transport system and the ammonium transporter AMT1 through direct protein-protein interaction (12).

A different mechanism of PII control has been identified in the process, how PII conformations modulate gene expression in response to different C/N-ratios. Here, the effect of PII is mediated through a small signalling mediator protein called PipX (PII-interacting protein X), which acts as a transcriptional co-activator of the global nitrogen control transcription factor NtcA. The latter controls a large regulon of at least 51 genes that are activated and 28 that are repressed (13). The mediator PipX swaps between PII and NtcA bound states, thereby either tuning down or stimulating, respectively, the activity of NtcA (6, 14). Partner swapping of PipX occurs in response to effector molecules 2-OG and the ATP/ADP balance (13).

In a previous PII-pull down study, several putative PII interactors of unknown function had been identified (12). The most prominent among these was the product of the *sll0944* gene, a member of the NtcA regulon (13). Here, we identify how the novel PII interactor Sll0944 transmits PII signals to a metabolic key enzyme. The *sll0944* gene product is annotated in Uniprot (https://www.uniprot.org/uniprot/P77971) as a 164 amino acid “uncharacterized protein” of unknown function. Close homologues are widespread in the cyanobacterial phylum according to BLAST analysis that points at an important function of this protein in the cyanobacterial metabolism. We previously encountered the *sll0944* gene during analysis of the nitrogen-starvation response of the model cyanobacterium *Synechocystis* PCC 6803. Both, a transcriptomic analysis (15) as well as a proteomic study (16) highlighted Sll0944 as highly enriched during long-term nitrogen-starvation induced chlorosis. Following nitrogen-depletion, the CO_2_ fixation products are redirected towards glycogen synthesis and concomitantly, the phycobiliproteins and the entire photosynthetic machinery is proteolytically degraded (15). At the same time, the carbon-polymer polyhydroxybutyrate (PHB) slowly accumulates in granular structures, which are derived from glycogen turn-over (17, 18). When the metabolic activities in the cells have reached a minimum, the cells reside in a dormant-like state, in which they can survive for months. As soon as a combined nitrogen source becomes available again, these cells rapidly awake and resume metabolism (15, 18, 19). At the same time, the levels of Sll0944 gradually returned to lower levels (19).

This study aimed to clarify the role of the Sll0944 protein in *Synechocystis* and its involvement in PII signalling. Our results show that Sll0944 regulates the glycolytic carbon flux at the phosphoglycerate mutase step in a PII-dependent manner through interaction with 2,3-phosphoglycerate-independent phosphoglycerate mutase (PGAM) in response to the nitrogen status. This establishes PGAM was a key control point of cyanobacterial carbon flow, as predicted previously by kinetic modelling of the cyanobacterial low carbon response (20, 21). Hence, we identified Sll0944 as PII-interacting protein that acts as a key regulator of PGAM in cyanobacterial carbon metabolism. We therefore named the *sll0944* product PirC (**P**II **i**nteracting **r**egulator of **c**arbon metabolism).

## Results

### In silico analysis reveals high conservation of Sll0944 (PirC) among cyanobacteria

According to the Uniprot database, the gene *sll0944* (from now on named PirC) from *Synechocystis* PCC 6803 codes for a 164 amino acid “uncharacterized protein” (https://www.uniprot.org/uniprot/P77971). A search for orthologues revealed that PirC is highly conserved among cyanobacteria, being present in almost all available genomes. All orthologs contain the Domain of Unknown Function (DUF) 1830. Further, a predicted NtcA-binding site 5’-GTN_10_AC-’3 (22), which is responsible for nitrogen-starvation induced expression, is found upstream of all homologous open reading frames, indicating conservation of nitrogen-starvation induced expression. Gene neighbouring analysis of 53 cyanobacteria reveals that in 67% genomes the *pirC* homologous are next to genes encoding radical SAM-like proteins, annotated as Elongator protein 3 (**Fig. S1**), which was recently shown to be a non-canonical tRNA acetyltransferase (23). In *Synechocystis* sp. PCC 6803, the *pirC* gene is upstream of *glgA1*, which encodes the major glycogen synthase that is required for acclimation to nitrogen deprivation (19).

Protein sequence comparisons showed that the first 52 amino acids at the N-terminus of PirC are not conserved in any of the other orthologues (**Fig. S2**), The high similarity to other PirC orthologs starts with methionine at position 53, which points to a wrong annotation of the translational start. In fact, the experimentally validated transcriptional start sites from *Synechocystis* PCC 6803 (24) gives rise to a shorter transcript with Met 53 as the most probable translational start site for PirC. Attempts to heterologously express the long and the short version in *E. coli* revealed only a correct folded and soluble protein of the short one. This finding agrees with the assumption that the 112 amino acids version of PirC being the physiologically relevant protein. Especially the 80 N-terminal amino acid residues are highly conserved within all cyanobacterial orthologs. Hence, all subsequent work was performed with the protein starting with the N-terminal sequence MSQ.

### PirC is a strong PII-binding partner

First experiments were set out to verify our previously suggested direct interaction between PII and PirC by an in-batch pulldown assay (Fig. S3 A). PirC co-eluted with PII in the presence of ATP or ADP but not in the presence of ATP plus 2-OG. The obstruction of PII interactions by 2-OG is typical for many PII interactions and indicates specificity (25). In the controls without PII, the elution fractions did not contain any PirC. To gain further insight into the nature of the PII-PirC complex, we used multi-angle light scattering coupled to size exclusion chromatography (SEC-MALS) to determine the size of the complex. First, the two proteins were analysed separately and then in a equimolar mixture (20 nmol monomers each), always in the presence of 5 mM MgCl_2_ and 0.5 mM ATP (**Fig. S3**). Most of the pure PirC eluted in a broad peak indicating formation of different oligomeric states. In addition, a small distinct peak appeared, for which MALS determined a molecular mass of 28.10 ± 0.07 kD (**Fig. S4 G**). This corresponds well to a dimer of recombinant PirC (calculated mass of monomeric recombinant PirC: 14.3 kDa). Under the same experimental conditions, pure recombinant PII protein eluted as a monodisperse species with a size of 45.7 ± 0.09 kDa, fitting perfectly to the trimeric structure of PII (calculated mass of trimeric recombinant PII: 46.9 kDa). When both proteins were mixed in a 1:1 molar ratio (monomers), two new peaks appeared with apparent molecular masses of 58.5 ± 0.2 kDa and 113.4 ± 2.5 kDa as determined by MALS. SDS-PAGE analysis showed that these two peaks indeed contained both proteins. Next, we first purified the PII-PirC complex using a two-step purification protocol, dialyzed it overnight against running buffer containing 2 mM ATP and then analysed it by SEC-MALS. As shown in Fig. S4 E, the chromatogram showed again the two molecular species with the dominant peak exhibiting an apparent molecular mass of 61.6 ± 0.1 kDa and a minor peak with an apparent molecular mass of 111.4 ± 2.2 kDa. Both peaks contained PirC and PII in the same ratio, with the protein band of PII being distinctly stronger than that of PirC. Taking the apparent mass of trimeric PII in the complex as fixed (45.7 kDa), the approx. 60 kDa complex most probably contained one PII trimer bound to monomeric PirC, whereas the larger complex most probably corresponds to a dimer of the smaller complex. The PirC-PII complexes seem to be very stable because no decay products of the pure complex were detected in the chromatogram.

### P_II_-PirC complex formation requires adenyl-nucleotides and responds to changing 2-OG levels

The effector molecule-dependence of the PII-PirC complex formation was quantitatively analysed by biolayer interferometry (BLI) spectrometry. C-terminally His_8_-tagged PII was bound to the Ni-NTA biosensor. Then, Strep-tagged-PirC at varying concentrations was allowed to bind to sensor-bound PII in a 180 sec association step followed by a dissociation step of 300 sec. A weak PII-PirC complex was formed without any effectors shown by the steady-state plot of binding and subsequent fast dissociation (**Fig. 1 A and B)**. In the presence of ATP or ADP, PirC-binding to PII strongly increased. Quantitative measurements in the presence of 2 mM ATP or ADP showed an apparent *K*_D_ values for PII-PirC complex formation of 37.3 ± 2.5 nM or 14.1 ± 0.7 nM, respectively. Similar results were obtained with SPR spectrometry. When the SPR sensor chip was loaded with 1000 RUs of His-tagged PII, the maximum response after injection of PirC in presence of ATP over the PII-loaded sensor was 270 RU (**Fig. S4 B**). As in SPR spectrometry the response signal in RUs is proportional to the mass change on the sensor, the mass increase of 270 RUs by PirC on 1000 RUs of PII-loaded sensor is close to one PirC monomer per PII trimer bound, agreeing with the suggested 3:1 monomeric stoichiometry from SEC-MALS analysis.

**Fig. 1.**
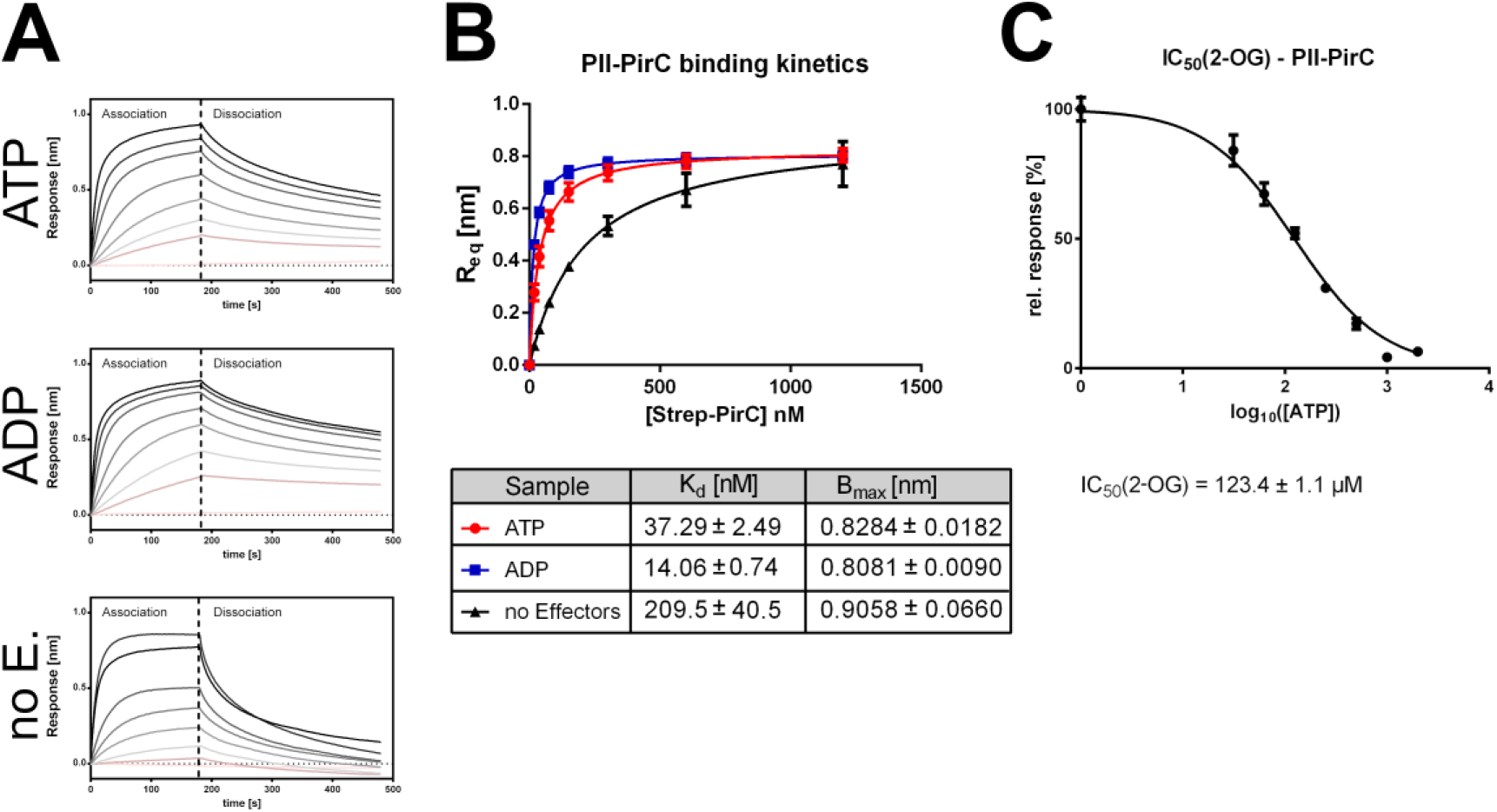
Complex formation of PII with PirC and modulation by effectors ADP, ATP and 2-OG. (A) *In vitro* binding studies by Bio-layer interferometry. His-tagged PII that was immobilized on sensor tips was allowed to associate for 180 s with PirC followed by 300 s dissociation. The overlay of experiments with increasing concentrations of PirC (9.375 nM – 1500 nM) is shown Plot of the binding responses in absence or presence of 2 mM ATP or 2 mM ADP form part (A) for the calculation of binding constants (depicted below) (C) Determination of apparent IC50 of 2-OG in presence of 2 mM ATP. All experiments were performed in triplicates and corresponding standard deviation are shown in B and C.

The inhibitory effect of 2-OG on PII-PirC interaction, observed in the initially performed pull-down experiments, was quantified by BLI spectrometry through a series of binding assays with increasing 2-OG concentrations in the presence of 2 mM ATP. The IC50 for 2-OG to completely prevent complex formation was determined as 123,4 ± 1.1 µM. This is close to the K_D_ of the third (lowest affinity) 2-OG binding site of PII (26)suggesting that PII with all three 2-OG sites occupied does not bind to PirC.

### Physiological role of PirC in Synechocystis

The above analysis revealed that PirC is a *bona-fide* PII interactor. The high conservation in the cyanobacterial phylum, including conservation of the NtcA-binding site, indicated an important function in nitrogen acclimation that is deeply rooted in cyanobacterial metabolism. To reveal such a function, we constructed a *pirC*-deficient mutant (Δ*pirC*) as well as two complementation strains (**Fig. S3**). One strain complemented with the native *pirC* gene (Δ*pirC*::*pirC)* and the second strain with a gene fusion, in which enhanced GFP (eGFP) was N-terminally fused to PirC under the control of the native *pirC* promoter (Δ*pirC*::*pirC-eGFP*). The latter strain allowed the subcellular localization of PirC by fluorescence microscopy (see below).

The wild type and mutant Δ*pirC* grew similarly well under non-stressed, nitrogen-sufficient conditions when exposed to day/night (12 h day and 12 h night) or constant light conditions (**Fig. S6 A**). After onset of nitrogen starvation under continuous light conditions, the Δ*pirC* mutant rapidly lost viability whereas the impact of nitrogen starvation was less severe under a day/night regime (12 h day and 12 h night) (Fig. S6). Hence long-term nitrogen chlorosis was investigated under permissive conditions in a day/night regime. The parameters growth (as increase in optical density, OD_750_), pigmentation, glycogen, and PHB content were recorded over one month. Overall, pigment degradation under day/night conditions required 21 days (**Fig. S7**) compared to only 5-7 days in continuous light (15). As inferred from photographs and absorbance spectra of cultures, pigment degradation was retarded in the Δ*pirC* mutant compared to the wild type and complemented strain. Likewise, the final stationary optical density (OD_750_) did not reach the same level in the mutant Δ*pirC* as compared to the wild type and the complemented strain (**Fig. 2 A**). Whereas the visual phenotype in chlorosis appeared rather modest, a striking difference between Δ*pirC* and the controls was observed with respect to glycogen accumulation. The wild type and complemented strain showed the typical rapid and steep increase in cellular glycogen levels, whereas the Δ*pirC* mutant showed only a minor increase in glycogen, reaching after two days only 28% of the wild type level and subsequently declined again. By contrast, the wild type and complemented strain Δ*pirC*::*pirC* maintained high glycogen levels over the course of chlorosis. As opposed to glycogen, the Δ*pirC* mutant accumulated significantly more PHB (up to 49% of the cell dry mass) than the wild type and the complemented mutant Δ*pirC*::*pirC* (30% or 29% PHB per cell dry mass, respectively) (**Fig. 2 B**). To confirm this result, PHB granules were stained with Nile red and visualized microscopically, as well as TEM micrographs were made from cells after 35 days of nitrogen depletion (**Fig. 2 C**). A much higher PHB content in the Δ*pirC* mutant than in the wild type or complemented strain could be clearly confirmed with these two PHB visualization techniques.

**Fig. 2.**
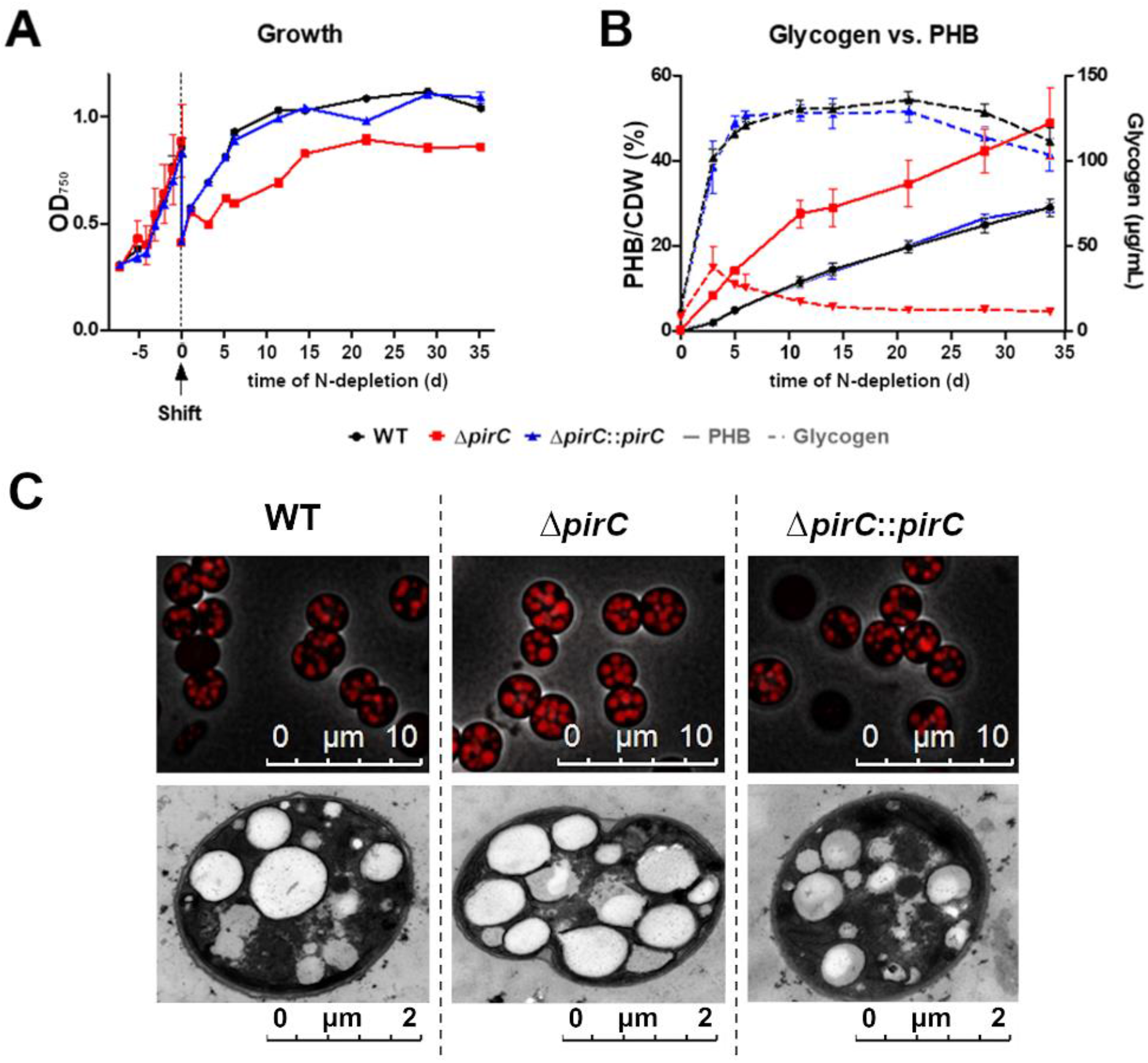
Effect of PirC deletion (Δ*pirC*) and its complementation (Δ*pirC::pirC*) on growth and carbon storage polymer accumulation during chlorosis. Graphs represent mean and SD from three biological replicates. (A) Growth curve, represented by OD_750_, seven days before nitrogen deficiency shift and during 35 days of chlorosis. (B) Glycogen and PHB content during nitrogen chlorosis. PHB content: plane lines; Glycogen content: dashed lines (C) Fluorescent micrographs and TEM of cells after 35 days of chlorosis. Upper row: 3D-deconvoluted picture of overlay of phase contrast- and fluorescence microscopy of Nile Red stained cells (1000 x magnification) Bottom row: TEM Pictures (5000 x magnification). Representative pictures of three biological replicates.

### Identification of the Sll0944-controlled process

The above studies pointed towards a crucial role of PirC in carbon storage metabolism during nitrogen starvation, a condition where the biochemical studies predicted release of PirC from PII. To find a mechanistic explanation for the disturbed carbon-storage phenotype of the PirC mutant, we aimed to identify further molecular targets of PirC. Therefore, co-immunoprecipitation (CoIP) experiments were conducted with crude extract from nitrogen-starved cells of an complemented strain Δ*pirC*::*pirC-mCitrine*. Preliminary experiments verified that the expression of PirC fused to mCitrine does also complement the mutant phenotype. CoIP experiments were performed with magnetic beads covered with an anti-GFP antibody and non-covered beads as a negative control. The pull-down was performed either in the presence of Mg^2+^, ATP and 2-OG, or in the absence of additionally supplemented effectors. The eluates from the independent experiments were analyzed after tryptic digestion by quantitative mass spectrometry. In the CoIP experiments performed in the absence of 2-OG, the only strongly enriched protein along with PirC was PII, conforming that PII is a major interaction partner at low 2-OG conditions (**Fig. S8**). The addition of 2-OG/ATP to the extract completely changed the CoIP pattern. In strong contrast, PII, was no more enriched, but the 2,3-bisphosphoglycerate-independent phosphoglycerate mutase (PGAM), encoded by the gene *slr1945*, appeared as dominant PirC interactor, which was significantly co-enriched together with PirC (**Fig. S9**). PGAM converts 3-phosphoglycerate (3-PGA) into 2-phosphoglycerate (2-PGA) in the beginning of lower glycolysis. In addition to PGAM, one further protein became also significantly enriched, CcmP (Slr0169), a putative carboxysome associated protein (27), however with a lower enrichment factor compared to PGAM.

### PirC is a competitive inhibitor of PGAM that is regulated by PII

The suggested interaction of PirC and PGAM is consistent with the lower glycogen and increased PHB phenotype of the Δ*pirC* mutant. This is because if PirC acts as inhibitor of PGAM, mutation of PirC should accelerate PGAM reaction, thus enhancing via glycogen catabolism the production of 2-PGA in lower glycolysis, which results in higher production of acetyl-CoA, the precursor metabolite of PHB. Therefore, we focused on verifying a putative negative regulation of PGAM by PirC. To validate the interaction of PirC with PGAM, the enzyme was prepared as recombinant protein with an N-terminal His_6_-tag. First, we checked the influence of PirC on the catalytic activity using a coupled enzyme assay, where the conversion of 3-PGA to 2-PGA was coupled to an excess of enolase, pyruvate kinase and lactate dehydrogenase, with the final oxidation of one molecule of NADH per molecule 2-PGA formed. The His-tag was removed by Thrombin cleavage to prevent unintended interaction of the tag in catalysis. To exclude any effects of PirC on the coupling enzymes, we tested the conversion of 2-PGA by the coupling enzymes in presence or absence of PirC, without noting any difference (Fig. S10). When PirC was added at increasing concentrations to the PGAM enzyme assay, a clear PirC-dependent inhibition of PGAM was observed. PirC inhibited PGAM by increasing the K_M_ for the substrate 3-PGA rather than lowering the v_max_, which corresponds to a competitive inhibition. At near equimolar concentration of PirC (200 nM) and PGAM (166 nM), the catalytic efficiency was reduced to one third In the excess of PirC, the catalytic activity of PGAM could be reduced more than 10-fold **(Fig. 3 A, Table 1)**.

**Table 1.** Kinetic constants of PGAM under varying concentrations of PirC and changes of constants by addition of PirC interacting molecules. Binding constants of PGAM/PirC complex with or without the presence of the substrates.

| Inhibition of PGAM by PirC at varying concentrations |  |  |  |  |
| --- | --- | --- | --- | --- |
| PirC, nM | $K_M$, mM | $v_{max}$, nkat $s^{-1} \cdot mg^{-1}$ | $k_{cat}$, $s^{-1}$ | $k_{cat} \cdot K_M^{-1}$, $s^{-1} \cdot M^{-1}$ |
| 0 | $0.1266 \pm 0.006$ | $47.54 \pm 0.52$ | $2,776 \pm 0,030$ | $21927,5 \pm 0.0$ |
| 200 | $0.3584 \pm 0.022$ | $44.92 \pm 0,75$ | $2,623 \pm 0,044$ | $7475.1 \pm 0.0$ |
| 400 | $0.7056 \pm 0.050$ | $46.56 \pm 1.09$ | $2,719 \pm 0,063$ | $3853.2 \pm 0.0$ |
| 600 | $1.376 \pm 0.102$ | $47.11 \pm 1.39$ | $2.751 \pm 0,081$ | $1999.2 \pm 3.8$ |
| 800 | $1.715 \pm 0.145$ | $43.27 \pm 4.10$ | $2.526 \pm 0,239$ | $1616.6 \pm 4.3$ |
| Antagonistic Effect of PII and its modulation by 2-OG (0.4 mM ATP, 600 nM PirC, 600 nM PII<sub>3</sub>) |  |  |  |  |
| Modulators of PGAM | $K_M$, mM | $v_{max}$, nkat $s^{-1} \cdot mg^{-1}$ | $k_{cat}$, $s^{-1}$ | $k_{cat} \cdot K_M^{-1}$, $s^{-1} \cdot M^{-1}$ |
| none | $0.120 \pm 0.013$ | $82.6 \pm 1.9$ | $4.96 \pm$ | $39083.3 \pm 0.6$ |
| PirC | $5.069 \pm 0.525$ | $116.6 \pm 7.3$ | $7.03 \pm 0.44$ | $1386.9 \pm 35.6$ |
| PirC/PII | $0.1508 \pm 0.030$ | $97.57 \pm 4.4$ | $5.89 \pm 0.27$ | $39058.4 \pm 0.9$ |
| PirC/PII + 0.123 mM 2-OG | $0.8543 \pm 0.501$ | $99.42 \pm 8.5$ | $5.33 \pm 0.52$ | $6239.0 \pm 4.6$ |
| PirC/PII + 1 mM 2-OG | $2.999 \pm 0.114$ | $88.45 \pm 4.1$ | $5.99 \pm 0.53$ | $1997.3 \pm 18.0$ |
| Binding constants for PGAM-PirC binding and modulation by substrates |  |  |  |  |
| Sample | $K_D$, nM | $IC_{50}$, mM | | |
| No Effectors | $2.89 \pm 34.49$ | - | | |
| 2-PGA | $1702 \pm 744$ (at $IC_{50}$) | $0.97 \pm 0.001$ | | |
| 3-PGA | $459.1 \pm 32.0$ (at $IC_{50}$) | $2.4 \pm 0.001$ | | |

**Fig. 3.**
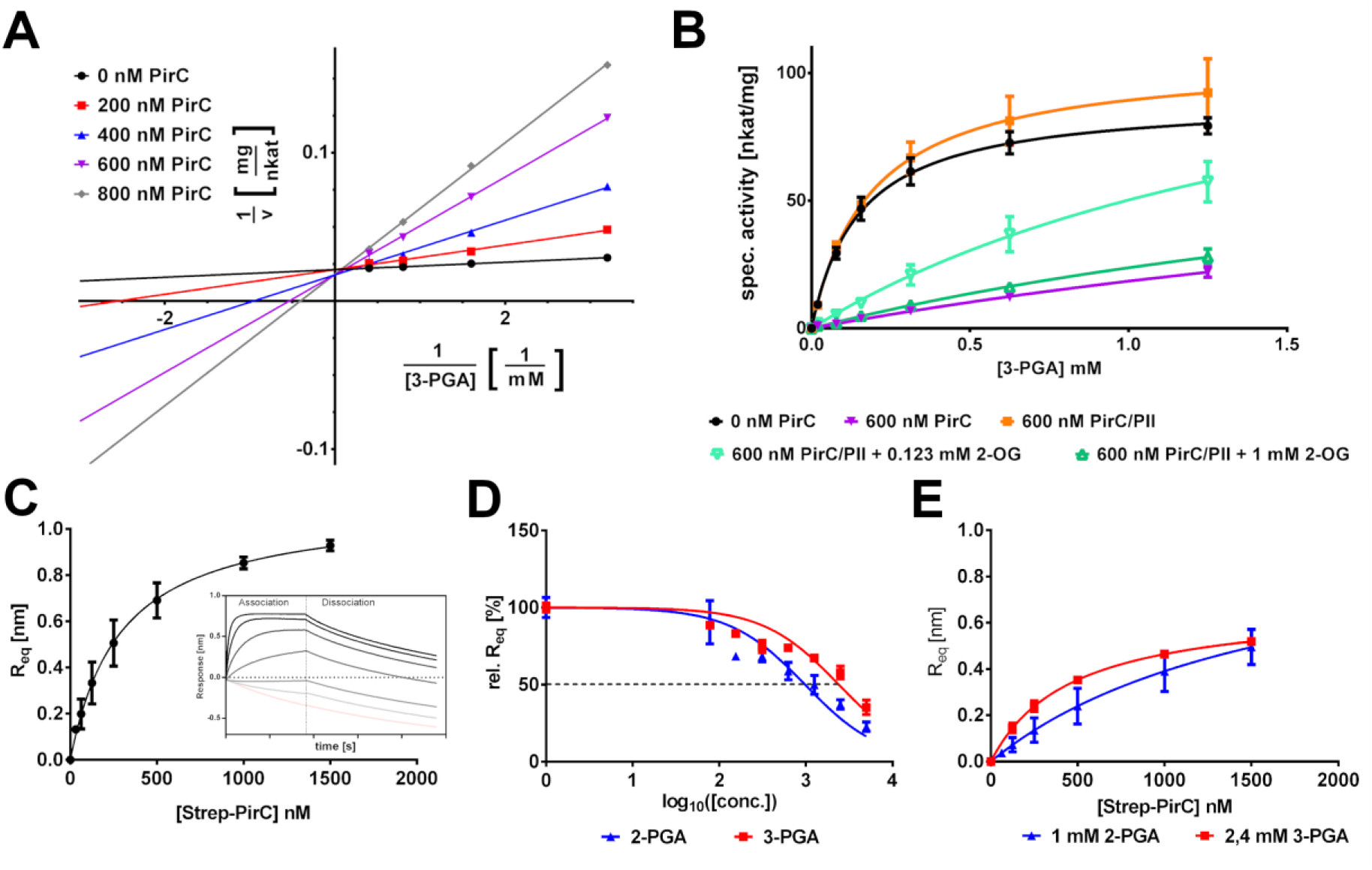
Effect of PirC and its modulation by PII and 2-OG on PGAM enzyme activity and PGAM-PirC complex formation in presence of 2-PGA and 3-PGA. (A) Lineweaver-Burk plot of PGAM specific activity at different PirC concentrations with the corresponding concentrations indicated. Each point represents the reciprocal calculation of the mean of triplicates. Control of PirC effect on coupled enzymes in shown in supplements (Fig. S6). Calculated constants are shown in Table 1. (B) Michaelis-Menten plot of the effect of PII on PGAM-PirC regulation in presence of 0.4 mM ATP and 2-OG at 0.123 mM and 1 mM. Each point represents the mean of triplicates. (C) Steady-state graph of PirC-PGAM binding assay using BLI. The mean of the R_eq_ value (three independent replicates) was plitted against the molar concentration of Strep-PirC. The error bars represents the SD. The inset shows the raw binding curves with different PirC concentrations. (D) Competetive inhibition of PirC-PGAM interaction by 2-PGA and 3-PGA. Plot for determination of IC_50_ (E) Steady-state graph of PirC-PGAM binding in presence of 2PG or 3PG at their IC50 concentrations.

The interaction between PII and PirC suggested that PII might regulate the inhibitory interaction of PirC with PGAM, in analogy to the effect of PII on PipX-NtcA interaction (14). To test this assumption, we performed PGAM assays that contained 600 nM PirC in presence of PII (600 nM trimeric concentration), supplemented with either ATP or ATP/2-OG. As shown in **Fig. 3 B**, addition of PII in the absence of 2-OG completely cancelled the inhibitory effect of PirC on PGAM activity and the K_M_ for 3-PGA returned to the value of non-inhibited PGAM **(Table 1)**. When, however, 1 mM 2-OG (corresponding to high carbon low nitrogen conditions) was added, PirC was again able to inhibit PGAM, like in the absence of PII. When 2-OG was added at a concentration of 0.123 mM (corresponding to the IC50_2-OG_ value of PII-PirC-complex formation) the inhibition of PGAM was approximately 50 % of the maximal inhibition by PirC alone. These results clearly show that *in vitro*, PII tunes the inhibition of PGAM by PirC in response to the 2-OG levels.

To further investigate the nature of PirC-PGAM binding, in particular with respect to the competitive mode of inhibition, Bio-layer Interferometry (BLI) assays were performed using His_6_-tagged PGAM bond to Ni-NTA-coated biosensor tips and Strep-tagged PirC protein as analyte. Strep-PirC at varying concentrations was allowed to bind to PII in a 180 sec association step followed by a dissociation step of 300 sec. A PGAM-PirC complex was formed without any effectors shown by the steady-state plot of the binding (**Fig. 3 C**). The competitive inhibition mode suggested that the substrates of PGAM should affect in turn the interaction of PGAM with PirC. Indeed, experiments in the presence of the PGAM substrates 3-PGA and 2-PGA (for the forward and backward reaction, respectively) revealed inhibitory effects on complex formation. With an IC_50_ of 0.97 mM, 2-PGA inhibited complex formation 2.4-times stronger than 3-PGA (IC50 = 2.4 mM) (**Fig. 3 D**). When the metabolites were assayed at their IC50 concentration (see above), 2-PGA (1 mM) increased the K_D_ to 1702 nM and 3-PGA (2,4, mM) to 459 nM, respectively (**Fig. 3 E, Table 1**).

### PirC deletion leads to accumulation of metabolites of lower glycolysis

The above-described analysis of the PGAM-PirC-PII triad demonstrated inhibition of PGAM activity by PirC, modulated by PII in response to 2-OG. This suggested that the inhibition of PGAM via PirC supports the formation of high glycogen levels during nitrogen starvation through diminished carbon catabolism via lower glycolysis in wild-type cells, while the absence of this inhibition resulted in higher glycogen catabolism via glycolysis in the mutant Δ*pirC*. To further verify this hypothesis, metabolite levels were analyzed from cells of the wild type and the Δ*pirC* mutant that were shifted from nitrate-containing into nitrogen-depleted (-N) medium. Samples were withdrawn after 0, 6, 24 and 48 hours for metabolome analysis. Nitrogen depletion had the expected effect on the total cellular steady-state metabolite pools, i.e. soluble amino acids were depleted to large extent, while organic acids accumulated. This switch resulted in lowered N/C ratios in cells under -N conditions. The decrease was faster in mutant Δ*pirC* than in the wild type. The fast decay of soluble amino acids in Δ*pirC* correlates with its slower bleaching, i.e. less amino acids are mobilized via pigment degradation than in wild type cells (Fig. S11). Most organic acids participating in the TCA cycle such as citrate, malate and succinate accumulated in both strains in a similar manner when shifted to –N conditions (**Fig. 4**). Also, the products of ammonium assimilation via GS/GOGAT, glutamine (Gln) and glutamate (Glu), showed similar changes in the wild type and Δ*pirC* with rapid decrease in Glu and slower decrease in Gln.

**Fig. 4.**
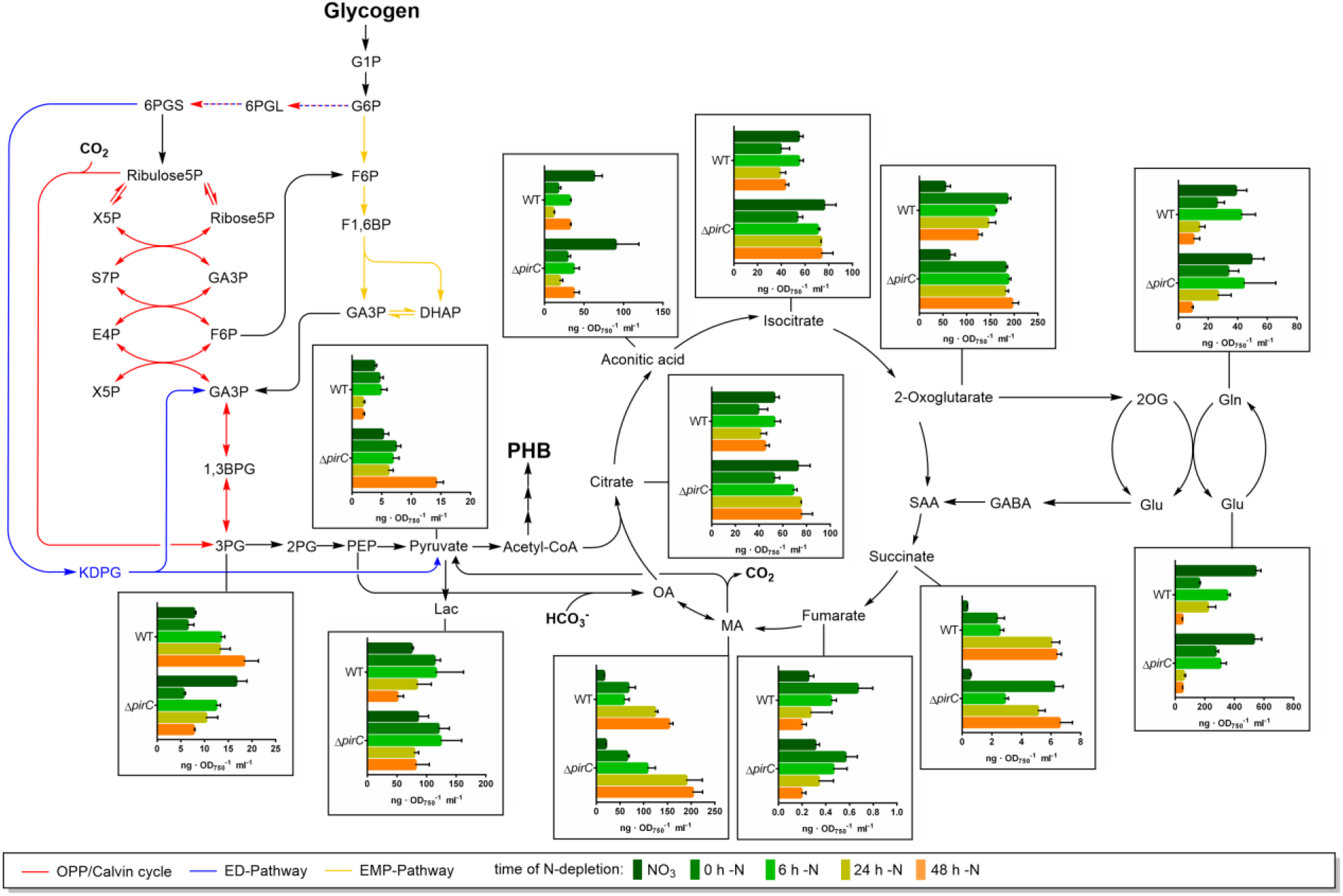
LC-MS analysis of general metabolite steady state levels during short term nitrogen starvation (48 h) of WT and Δ*PirC* depict in metabolic background. Each bar represents two independent biological replicates with each in technical duplicates. The error bars represent the SD of the combined data. The values are in ng · OD_750_ ^−1^· ml^−1^. Result of the hole metabolite analysis is shown in Fig S11.

Besides these general metabolic responses, where the Δ*pirC* mutant showed no discernable differences in metabolite pools, a few very specific and intriguing differences were recorded at decisive steps: The N-status reporter molecule 2-OG accumulated immediately after the shift to N-free medium in both strains. In the wild type the 2-OG levels decreased gradually over the following 48 h, whereas they remained constantly elevated in Δ*pirC* cells, they even increased slightly (**Fig. 4**). Even more remarkable, the 3-PGA concentration increased in the wild type over the course of N-starvation, whereas it gradually declined in the Δ*pirC* mutant. The increasing 3-PGA levels, substrate of the PGAM, in the wild type cells are indicative of an inhibition of the PGAM reaction. Lack of PGAM inhibition by PirC in the mutant explains their lower 3-PGA levels. 3-PGA is known as an allosteric activator of the glucose-1-phosphate-adenylyltransferase (*glgC*) in bacteria, which catalyzes the initial step of the glycogen synthesis (28). Hence, the lower 3-PGA levels in the Δ*pirC* mutant are also consistent with its lower glycogen accumulation (see **Fig. 2**). Hence, stimulated glycogen accumulation in N-starved wild type cells is supported by the PirC blockage of lower glycolysis after its release from PII and an allosteric stimulation of GlgC. Downstream of the PGAM reaction, the levels of pyruvate could be detected. They responded inversely to 3-PGA, with decreasing levels in the wild type but a strong increase in the Δ*pirC* mutant. After 48 h of N-starvation, the pyruvate level in the Δ*pirC* mutant was 14-fold higher than in the wild type. Again, this observation is consistent with an increased flux through the PGAM reaction due to the missing inhibition by PirC, since the produced 2-PGA is further converted into pyruvate. The higher carbon flux through lower glycolysis result in higher pyruvate levels in the Δ*pirC* mutant, thereby lowering the opposite carbon flux into glycogen and are also consistent with its increased PHB levels, which are derived from acetyl-CoA, the immediate reaction product from pyruvate.

### Subcellular localization of PirC

To finally reveal subcellular localization of PirC in different growth stages, where it presumably interacts preferentially with either PII or PGAM (as deduced from our *in vitro* studies), we microscopically analysed the fluorescence signal from cells of the Δ*pirC*::*pirC-eGFP* strain. During exponential growth in nitrate-containing BG_11_ medium (predicted PII-PirC complex), the eGFP signal was centrally localized in the cytoplasm (**Fig. S12**). After shifting the cells to nitrogen-depleted medium, increasing 2-OG levels (see metabolite levels above) should promote dissociation of the PII-PirC complexes and instead, allow PirC-PGAM interaction. In fact, the localization of PirC-eGFP changed after nitrogen-downshift, such that during the first 24 h of nitrogen starvation, the centrally localized eGFP signal slowly expanded to the peripheral region of the cytoplasm where it then remained throughout chlorosis. This result corroborated the dynamics of PirC interactions and its response to nitrogen limitation.

## Discussion

In this work, we identified a novel key control point of cyanobacterial carbon metabolism, the glycolytic phosphoglycerate mutase (PGAM) reaction, converting 3-PGA into 2-PGA. In animal systems, where glycolysis provides core reactions for energy supply, it has been shown that glycolytic breakdown of glucose-6P is mainly regulated at the phosphofructokinase step according to the energy demand of the cells (29).By contrast, in photoautotrophic organisms, glycolytic steps are used in both directions, in gluconeogenetic direction towards glycogen or starch synthesis, and in the opposite, glucose catabolic direction, to produce precursors for multiple biosynthetic routes required for cell growth. In this regard it is important to note that the PGAM reaction is at the branch point of newly fixed CO_2_: 3-PGA, the first stable reaction product from RuBisCO-catalyzed CO_2_ fixation, can go in two different directions: most of it is converted into 2,3-bisphosphoglycerate (2,3-PGA) and further to glyceraldehyde-3-phosphate (G3P), from which the acceptor of RuBisCO, ribulose 1,5-bisphosphate (Ru1,5-BP), is regenerated via the Calvin-Benson cycle reactions.. Part of G3P can be converted to the main storage compound glycogen (in plants, starch) via gluconeogenic reactions. Alternatively, 3-PGA can exit the Calvin-Benson cycle by its direct conversion via PGAM to 2-PGA, which is further metabolized in lower glycolytic reactions. From there, the majority of cellular amino acids and lipids are derived in photoautotrophs, with pyruvate, acetyl-CoA and 2-OG representing key metabolites.

The PGAM reaction as a key control point of carbon metabolism has been predicted previously by kinetic modelling of the cyanobacterial low carbon response (20, 21). It has been shown that 2-PGA accumulates to high amounts (5-7 times) in cells shifted from high CO_2_ (5%) to ambient air (0.04% CO_2_) in *Synechocystis* sp. PCC 6803 (30) as well as in *Synechococcus elongatus* PCC 7942 (31). The high 2-PGA accumulation was taken as indication that under carbon-limiting conditions more freshly fixed organic carbon is deviated by the PGAM reaction from the Calvin-Benson cycle into lower glycolysis to sustain biosynthesis of amino acids and other cellular compounds. Here we provide *in vitro* and *in vivo* evidences, that for acclimation to nitrogen starvation, the PGAM reaction, catalysed by the *slr1945* product, represents a key control point. This control operates through a novel regulatory mechanism, in which the small regulatory protein PirC acts as a mediator of the signal from the pervasive PII regulatory protein to tune the activity of PGAM, a control mechanism so far not known for any enzymatic reaction. To further understand the competitive inhibition of PGAM by PirC, as demonstrated here through kinetic and binding studies, structural analysis of the enzyme complexes is required.

According to our model depicted in **Fig. 5**, under nitrogen-sufficient conditions, where 2-OG levels are low, PirC is bound by PII, preventing the interaction with PGAM. Efficient conversion of 3-PGA to 2-PGA directs newly fixed carbon towards lower glycolysis to support the synthesis of amino acids and fatty acids. Only a minor fraction is converted into glycogen. When the cells experience nitrogen limitation (causing increased intracellular 2-OG levels), the PII-PirC complex dissociates and PirC instead interacts with PGAM, thereby inhibiting its enzymatic activity. The nitrogen down-shift goes along with a re-localization of the PirC-eGFP signal, indicating localization of the PirC-PGAM complexes. Accordingly, PGAM would be located throughout the entire cytoplasmic space of the cells. By contrast, PirC in complex with PII localizes to the central region of the cell. This agrees with the preferential localization of PII in nitrate-grown cells, reported by Watzer et al. (11). As a consequence of the PirC-PGAM interaction, conversion of 3-PGA to 2-PGA is blocked, leading to increased 3-PGA levels, which is now re-directed towards the accumulation of glycogen. Due to this metabolic switch, the flux towards amino acid synthesis is slowed down, adjusting these pathways to the limited supply of nitrogen. Furthermore, 3-PGA is an allosteric activator of GlgC, the glucose-1-phosphat-adenylyltransferase (GlgC), which catalyzes the initial and regulated step of the glycogen synthesis (28). Hence, the PirC mediated PGAM inhibition not only slows down lower glycolysis but is also stimulating glycogen accumulation via the enhanced GlgC activation. The glycogen levels increase, until the cells are densely packed with glycogen, which can amount up to 50 % of the cell dry weight (15). Already after 24 hours of nitrogen starvation, the glycogen stores can be filled, and they remain at high levels throughout chlorosis. Recent data indicate a constant turn-over of glycogen until the cells enter complete dormancy after about 7-10 days of N-starvation (17). In *Synechocystis*, which expresses the PHB synthesis machinery, the acetyl-CoA molecules arising from the remaining glycolytic flux are directed towards PHB synthesis under these non-growing conditions. The amount of PHB steadily increases during prolonged nitrogen starvation. In the PirC mutant, PGAM cannot be appropriately inhibited. Therefore, increased flux towards 2-PGA and lower glycolysis leads to massive over-accumulation of PHB. In agreement with this model, the PII-deficient *Synechocystis* mutants are unable to accumulate PHB during nitrogen chlorosis (32). In the absence of PII, PirC will constantly inhibit PGAM activity, explaining decreased metabolite levels downstream of 2-PGA (10), limiting the supply of acetyl-CoA for PHB synthesis.

**Fig. 5.**
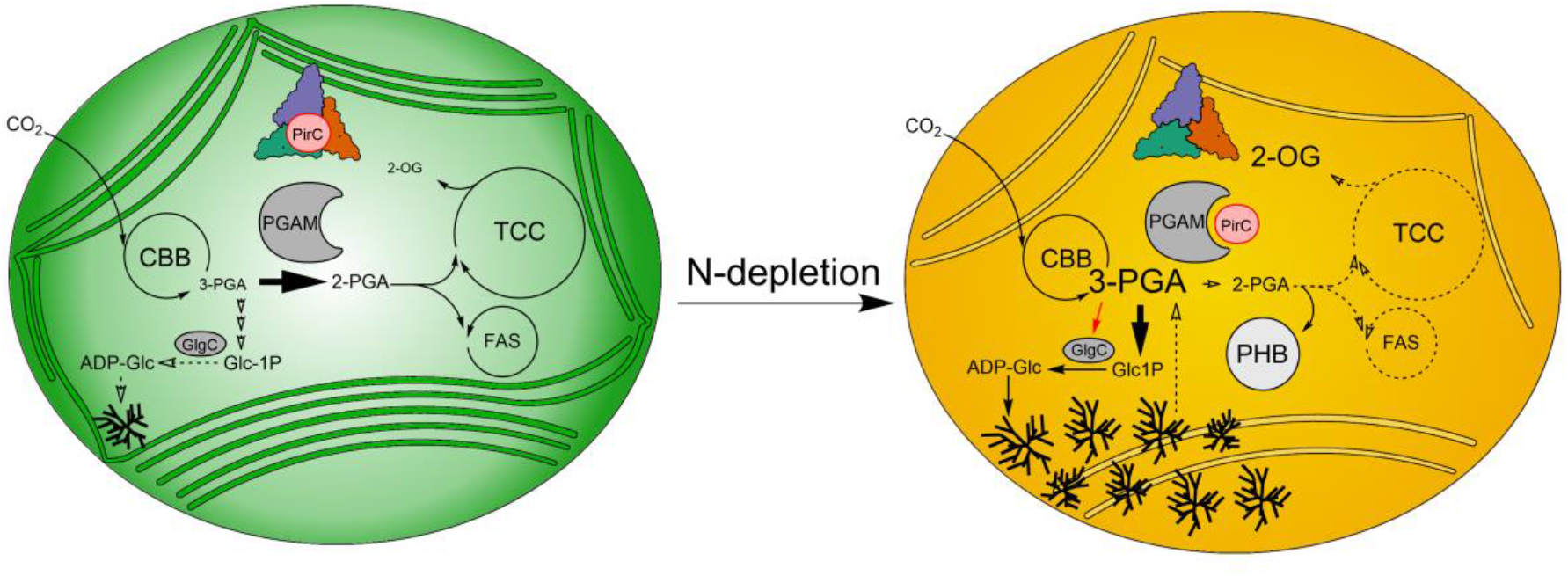
Model of regulation of central carbon metabolism by PirC, PII and PGAM interactions. In vegetative cells, when 2-OG levels are low, PII binds to PirC and prevents the inhibition of the PGAM. PGAM directs 3-PGA downstream to the biosynthesis of fatty acids (FAS) and amino acids (TCC). However, when the cells become N-depleted, 2-OG levels increase and promote release of PirC from PII. PirC inhibits the PGAM followed by an elevation of the 3-PGA concentration. 3-PGA enhances the activity of GlgC and directs the carbon flux equilibrium back to glycogen, resulting in enhanced accumulation of glycogen and an attenuated catabolism of glycogen during chlorosis. During chlorosis. acetyl-CoA, formed from glucose catabolism, is converted into PHB.

This novel glycolytic switch at the PGAM enzymatic step via the small PII-interacting regulatory protein PirC is reminiscent to the control of transcription by NtcA via the small PII-interacting protein PipX. The latter is either complexed by PII under low 2-OG conditions or bound to NtcA at elevated 2-OG levels (6). Both small PII-mediator proteins functionally interact to coherently reprogram metabolism and gene expression under low nitrogen conditions. Through release of the PII-PirC complex in response to increasing 2-OG levels, PirC tunes down PGAM. This response is further amplified by the concomitant dissociation of the PII-PipX complex, where PipX co-activates NtcA-dependent expression of many low nitrogen-induced genes, among them the *pirC* gene. In chlorotic cells, *pirC (sll0944*) belongs to the most strongly up-regulated genes of the entire transcriptome (15),and in agreement, PirC is one of the most highly enriched proteins during chlorosis (16). In an advanced stage of chlorosis, this strong accumulation of PirC ensures tight inhibition of glycolysis, to maintain high glycogen levels, which are required for efficient resuscitation from chlorosis (19)

Through engagement of these mediator proteins, PII can largely expand its regulatory space. In principle, any activity, which can be modulated through interaction with a peptide, could be tuned by PII through fusing it to a PII interaction surface, giving rise to a new toolbox for synthetic biology. It is also reasonable to speculate that disabling the regulation of PirC can be used for metabolic engineering in cyanobacteria, in particular, for the bioproduction of metabolites derived from lower glycolysis, such as succinate, malate, or lipids as well as fatty acids. Furthermore, cyanobacteria provide the native potential for synthesis of isoprenoids or terpenes whose precursors are pyruvate and GAP. Some terpenes are promising compounds in cancer therapy and inhibit essential enzymes in tumor formation in a variety of cancer types (33–35). In addition, genetically engineered cyanobacteria can be used for the synthesis of heterologous compounds such as n-butanol. In previous attempts to generate a *Synechocystis* strain for the production n-butanol, production was limited by the supply of precursor molecules (36). Mutation of PirC could overcome these limitations. As a proof of principle, we have transformed the PirC-deficient mutant with genes for PHB synthesis from *Cupriavidus necator*, resulting in a strain which accumulates more than 80% PHB of cell dry mass, which is by far the highest accumulation of PHB ever reported in a cyanobacterium. This demonstrates impressively the significance and biotechnological potential of decoupling the regulation by PirC.

## Material and Methods

### Strains and cultivation

A list of all used strains for this study is provided in Table S1.

For preculturing and growth experiments, the *Synechocystis* sp. PCC 6803 (from now on *Synechocystis*) strains were cultivated in BG_11_ medium according to Rippka (37). The cultivations were performed in Erlenmeyer flasks without baffles whereby 50 ml cultures were cultivated in 200 ml flask and 200 ml cultures in 500 ml flasks. Typical cultivation was performed either at continuous illumination (∼50 μE m^−2^ s^−1^) or light-dark conditions (12 h light and 12 h darkness) and 28 °C while the cultures were shacked continuously at 125 rpm. For nitrogen depletion, cultures were cultivated in BG11 media without 17,65 mM NaNO_3_. Whenever necessary, appropriate antibiotics were added to the different strains to ensure the continuity of the mutation.

For nitrogen deficiency experiments, pre-cultures of *Synechocystis* were cultivated for three days as described previous at an initial OD_750_ of 0.1. Experimental cultures were then prepared in BG11 medium with a set starting OD_750_ of 0.2 and grown for two days under identical conditions until they reached an OD_750_ of 0.6-0.8. For the nitrogen shift experiments, cells from the cultures were harvested by centrifugation (4000 g, 10 min), washed with and resuspended in BG11_0_ medium to create cultures with an initial OD_750_ of 0.4.

Viability assays of *Synechocystis* were performed on BG_11_ agar plates (1,5 % agar) containing 20 mM TES buffer (pH 8.2) and 3 mM Na_2_S_2_O_3_. Cultures of an OD_750_ of 0.4 were diluted from 10^−1^ up to 10^−4^. Of each dilution, 5 μl were dropped on a BG_11_ agar plate. The plates were cultivated seven days at either continuous light or light-dark conditions.

Cultivation of *Escherichia coli* cultures was performed with LB medium and agar. Lennox broth: 5 g·l^−1^ Yeast extract, 10 g·l^−1^ Tryptone, NaCl 5 g·l^−1^, solid: 15 g·l^−1^ agar

### Plasmids and cloning

A list of all used primer, plasmids and its cloning procedure are listed in Table S2 and Table S2 as well as in Figure S5.

### Overexpression and Purification of Proteins

*Escherichia coli* Lemo21(DE3) were used for overexpression of the various kind of proteins. The expression of His-tagged proteins was performed as described in the manufactured expression protocol in 2-fold concentrated LB media. An overnight expression, induced by addition of 400 µM IPTG, in a 1 l culture was performed with 1 mM L-rhamnose at 25 °C during continuous shaking at 120 rpm. Additionally, dependent on the plasmid the appropriated amount of the antibiotic was added to the culture. The expression of Strep-tagged proteins based on pASK-Iba5Plus expression plasmid was induced by addition of 200 µg·l^−1^ anhydrotetracycline without addition of L-rhamnose because of the T7 RNA polymerase independent expression.

The heterologous proteins containing His-tags were purified via 1 ml Ni-NTA HisTrap columns (GE Healthcare). The cells were lysed in 50 ml lysis buffer containing 50 mM Na-phosphate buffer pH 8, 300 ml NaCl, 1 mM DTT, 1 mM Benzamidine and 0,2 mM PMSF. The His-tagged proteins were loaded on the Ni-NTA column with Buffer A containing 50 mM Na-phosphate pH 8, 300 ml NaCl and eluted via a gradient of increasing imidazole (0-500 mM, Buffer B) using a ÄKTAPurifier™ System (GE Healthcare). After this first purification, the proteins were further purified via size exclusion chromatography using a Superdex™ 200 Increase 10/300 GL (GE Healthcare) with 50 mM Tris/HCl buffer containing 100 mM KCl and 0.5 mM EDTA.

For purification of Strep-tagged proteins, 5 ml Strep-tactin® superflow columns were used. Cells were lysed in lysis buffer containing 100 mM Tris/HCl pH 8, 150 mM NaCl, 1 mM EDTA and 1mM PMSF. The proteins were loaded on the column and eluted with buffer containing 5 mM Desthiobiotin. The buffer of each purified protein was exchanged via dialysis using dialysis buffer (50 mM Tris/HCl pH 7.8-8, 100 mm KCl, 5 mM MgCl_2_, 0,5 mM EDTA, 40 % glycerol) and a 3 kDa cutoff dialysis tube. All purification steps were checked via SDS-PAGE according to previous studies (38).

The His-tag of PGAM was removed by Thrombin cleavage using Thrombin of bovine of Sigma Aldrich according to protocol (39).

### In-batch Pulldown assay

Interactions between PirC and PII were first checked via an in-batch pulldown experiment. For this, His_6_-PirC and Strep-PII were mixed in equimolar amounts and incubated for 20 min at room temperature in buffer containing 50 mM Tris/HCl, pH 7.8, 100 mM KCl, 0.5 mM EDTA, 10 mM MgCl_2_ and additionally either 2 mM ATP, ADP or ATP and 2-OG. The mixtures were applied to Strep-Tactin XT coated magnetic beads. After 30 min of incubation the magnetic beads were removed via a strong magnet, followed by three times washing step with incubation buffer. For analysis of bound proteins via SDS-PAGE, the beads were boiled 10 min at 100 °C as described previously (38).

### Size exclusion chromatography coupled with multiple light scattering (SEC-MALS)

Oligomeric state analysis and the size of protein complexes was analyzed by size exclusion chromatography coupled to multi-angle light scattering (SEC-MALS). This was performed using an purifier system connected to a Superose 6 Increase 10/300 GL column (both from GE healthcare, Solingen, Germany) at a flow rate of 0.5 ml · min^−1^ in running buffer (50 mM Tris/HCl (pH 8.0), 150 mM NaCl, and 10 mM MgCl_2_). The ÄKTA micro was connected to downstream MALS using the miniDAWN TREOS combined with an Optilab T-rEX (both from Wyatt Technology, Dernbach, Germany) refractometer. Data analysis was performed using the software ASTRA 7 (Wyatt Technology, Dernbach, Germany) and Unicorn 5.20 (Build 500) (General Electric Company, Boston, USA). The column was calibrated using the gel filtration calibration kit LMW and HMW (GE Healthcare, Solingen, Germany) according to the manufacturer’s instructions.

### Co-Immunoprecipitation and Liquid chromatography-Mass spectrometry (LC-MS/MS)

To identify putative interaction partners of PirC, co-immunoprecipitation experiments were performed. For this, *Synechocystis* ΔPirC::PirC-mCitrine cultures were pre-cultivated in100 ml BG11 medium to an OD_750_ of ∼0,8 and subsequently shifted to N-depleted medium. For Co-IP experiments in presence of the PII effector molecules, cells were harvested after 24 h N-depletion by centrifugation for 10 min at 4200 x g at 4 °C, followed by resuspension of the cell pellet in 2 ml binding buffer containing 100 mM TRIS (pH 7.5), 100 mM KCl, 1 mM MgCl_2_, 1 mM DTT, 0.5 mM EDTA, 2 mM ATP and 2-OG. The cells were lysed with 150 µl glass beads in 1.5 ml screw cap tubes by harsh shaking in a high-speed homogenizer for 5 times 30 sec shaking at speed of 7 m · s^−1^ with each 5 min break. The lysate was than centrifuged at 25,000 x g for 5 min at 4 °C and a supernatant volume corresponding to a protein yield of approx. 3 mg was used for the immunoprecipitation. Therefore, GFP-Trap Magnetic Agarose beads or control beads without antibodies were used according to the manufactured protocol (both Chromotek, Planegg-Martinsried, Germany). The loaded magnetic beads were heated for 10 min at 95 °C in SDS loading buffer for the dissociation of purified proteins. Protein solutions were subjected to short SDS-PAGE runs, in which proteins were allowed to migrate for 1.5 cm into 12% Bis-Tris Gels (Invitrogen) and then stained with Coomassie blue. Protein containing gel regions were isolated and subjected to InGel digestion with trypsin as described elsewhere (40). Tree independent experiments, each including a PirC-mCitrine and a control CoIP were performed in total. Peptides were subjected to a clean-up step using StageTips (41) and subsequently analyzed by mass spectrometry. LC-MS/MS analysis was performed on a Q Exactive HF mass spectrometer (ThermoFisher, Germany), using linear, segmented 60 min nanoLC RP gradients as described elsewhere (16). All raw data was processed using MaxQuant software suite (version 1.6.5.0) at default settings. MS2 peak lists were searched against a target-decoy database of the *Synechocystis* sp. PCC 6803 proteome, including the sequence of PirC(Sll0944)-mCitrine. Label free quantification was used to calculate LFQ intensities for each CoIP sample. Data from all experiments was analyzed via the Perseus software (version 1.6.5.0). For the identification of significantly enriched proteins in PirC-mCitrine CoIPs, a *t*-test was performed with the following requirements: each protein had to be detected in at least two replicates and an FDR of 0.01 at S0 = 0.1 was set.

### Biolayer interferometry using the Octet K2 system

*In vitro* binding studies were done by Bio-layer interferometry (BLI) using Octet K2 system (FortéBio). The experiments were performed in HEPES buffer (20 mM HEPES-KOH pH 8.0, 5 mM MgCl_2,_ 0.005 % NP-40, For protein interactions of His_8_-PII–strep-PirC 150 mM KCl and for the His_6_-PGAM–strep-PirC interaction 10 mM MnCl_2_ was added to the buffer. In the first step PII-His_8_ (400 nM, trimeric) or PGAM-His_6_ (500 nM) were immobilized on Ni-NTA sensors (FortéBio) followed by a 60 sec baseline measurement. For the binding of PirC, the biosensors were dipped into the PirC solution for 180 sec (Association), with concentrations ranged between 9.375 nM – 1500 nM. Dependent on the experiment different effector molecules were added to the binding buffer, ADP, ATP and 2-OG in PII binding studies, and 2PG as well as 3 PGA in PGAM binding assays. The assay was terminated by a 300 sec dissociation step. To prevent false positive results in each experimental set one measurement without any interaction partner was performed. The biosensors were regenerated after each use with 10 mM glycine (pH 1.7) and 10 mM NiCl_2_ as proposed in manufacturers recommendations. The recorded curves of a set were preprocessed by aligning to the average of the baseline step and to the dissociation step. The response in equilibrium (R_eq_) was calculated using the Data Analysis Software of the Octet System. The Concentration versus R_eq_ plots were made for each set of experiments which were then used to calculate the Dissociation constant K_D_.

### Glycogen measurement

The determination of the glycogen content of *Synechocystis* cultures was performed according to previous studies (15, 42). The glycogen was isolated from cell pellets of 2 ml culture. The glycogen was hydrolyzed to glucose with 4,4 U · µl^−1^ amyloglucosidase from *Aspergillus niger* (Sigma Aldrich) for 2 h at 60 °C. The resulting glucose concentration was measured via o-toluidine assay (43). The samples were boiled in 1:6 dilution with a 6 % o-toluidine reagent (in glacial acetic acid) for 10 min, then cooled on ice and measured at 635 nm. The concentration of samples was calculated using a calibration curve of defined quantity of glucose (0, 10 µg, 50 µg, 100 µg, 250 µg, and 500 µg).

### PHB Quantification

Polyhydroxybutyrate was detected by high-performance liquid chromatography as described previously (44). Eleven ml of chlorotic cultures were centrifuged at 4200 x g for 10 min and the cell pellet was vacuum dried in pre-balanced 2 ml reaction tubes. To calculate the cell dry weight (CDW) the tube was cradled again. Then the pellet was boiled for 60 min in concentrated H_2_SO_4_ (18 mol · l^−1^). Thereby, the cells were lysed and PHB converted to crotonic acid. Next, 110 μl of this solution was diluted 1:10 with 14 mM sulfuric acid solution and centrifuged 5 min at 25,000 × *g* followed by another 1:2 dilution. After another centrifugation step 300 μl of the clear supernatant was used for analytical HPLC. Reversed-phase HPLC was performed using the Chromatography system HP1090 M, equipped with a thermostated autosampler and diode-array-detector, HP Kayak XM 600 workstation. The crotonic acid was detected by measuring the absorbance at 210 nm. The crotonic acid concentration of the samples was calculated using a calibration curve of defined concentrations (0.5 mg·ml^−1^, 0.25 mg·ml^−1^, 0.125 mg·ml^−1^ and 0.0625 mg·ml^−1^)

### PGAM enzymatic assay

The PGAM activity and the effect of PirC was determined by a coupled enzyme assay as described previous (45, 46). For that, 10 µg of purified PGAM was used in a 1 ml reaction. The reaction mixture containing 20 mM HEPES-KOH (pH 8,0), 100 mM KCl, 5 mM MgSO_4_, 0.4 mM MnCl_2_, 50 µg·ml^−1^BSA, 1 mM DTT, 0.4 mM ADP, 0.2 mM NADH, 0.5 U enolase (Sigma Aldrich), 2 U Pyruvate kinase (Sigma Aldrich), 2 U Lactate dehydrogenase (Roche) and 10 µg PGAM was pre-warmed to 30 °C. The Assay was started by adding the 3-PGA solutions. The resulted decrease of NADH over time was recorded with Specord50 (Jena Analytics) at 340 nm. A blank assay without 3-PGA was also performed, no decrease was detectable.

### Phase contrast and fluorescence microscopy

The visualization of PHB granules was done by phase contrast fluorescence microscopy using the Leica DM5500 B with the Leica CTR 5500 illuminator. The integrated camera Leica DFC 360 FX was used for image acquisition. The settings were adjusted by the Leica Application Suite Advanced Fluorescence (LAS 4.0). The specimen was prepared by dropping 10 µl of cell culture on an agarose coated microscope slide and observed using the Leica HCX PL FLUATAR (100⨯ 1,30 PH3) with immersion oil for a 1000-fold magnification. The visual detection of PHB via fluorescence was performed by staining the cells with 0,33 µg·ml^−1^ Nile Red. The fluorescence of Nile Red was excited with light between x – y nm (CYR channel) and of GFP like proteins with light between 455-495 nm (GFP channel). The images were processed by 3D deconvolution of all channels using LAS.

### Transmission electron microscopy

For electron microscopic pictures, *Synechocystis* cells were fixed with glutaraldehyde and post-fixed with potassium permanganate, respectively. Afterwards, microtome sections were stained with lead citrate and uranyl acetate (47). The samples were then examined using a Philips Tecnai 10 electron microscope at 80 kHz.

### Metabolome analysis

For metabolome analysis by LC-MS *Synechocystis* was cultivated in 200 ml under N-depletion as described previously for 48 h under continuous lightning. The sampling was carried out 0, 6, 24 and 48 hours after the shift. Samples of 5 mL liquid culture were quickly harvested onto nitrocellulose membrane filters (ø 25 mm, 0.45 µm pore size, Porafil NC, Macherey-Nagel) by vacuum filtration, put in 2 ml Eppendorf microtubes, and immediately frozen in liquid nitrogen. Cells on filters were stored at −80 °C until analysis.

Extraction was done using LC-MS grade chemicals. To every filter, 630 µL methanol and 1 µL carnitine as internal standard (1 mg·ml^−1^) were added. Cells were re-suspended by rough mixing and then incubating samples in a sonication bath for 10 min. Samples were shaken for 15 min prior to addition of 400 µL chloroform and incubation at 37 °C for 10 min. Next, 800 µL of ultrapure water were added. The extracts were shaken for 15 min and then incubated at −20 °C for at least 2 h. Cell debris and filters were removed by centrifugation (20000 g, 5 min, 4 °C). The upper polar phase was transferred completely into a new microtube and subsequently dried by vacuum concentration (Concentrator plus, Eppendorf). The dried extracts were re-suspended in 200 µL deionized water and filtrated (0.2 µm filters, Omnifix-F, Braun). The filtrates were then analysed via the high-performance liquid chromatograph mass spectrometer LCMS-8050 system (Shimadzu), as previously described by Selim et al., 2018 (48). LC-MS data analysis was done using the Lab solution software package (Shimadzu).

## Data availability

Proteome raw data files acquired by mass spectrometry were deposited at the ProteomeXchange Consortium (http://proteomecentral.proteomexchange.org) via the PRIDE partner repository (48) under the identifier PXD021415. The pre-published dataset is password protected and available upon request to the authors.

## Acknowledgments

The project was funded by grants from the German Research Foundation Fo195/9-2 and as part of the research group (FOR2816) SCyCode “The Autotrophy-Heterotrophy Switch in Cyanobacteria: Coherent Decision-Making at Multiple Regulatory Layers” to MH (HA 2002/23-1) and BM (MA 4918/4-1). LC-MS/MS systems at the Department of Quantitative Proteomics were supported through the German Research Foundation (DFG) grants No. INST 37/935-1 and INST 37/741-1 FUGG. The LC-MS/MS equipment at University of Rostock was financed through the HBFG program (GZ: INST 264/125-1 FUGG). We want to acknowledge the contribution of initial studies on Sll0944 by Alexander Klotz that paved the way for the present study.

## Author’s contributions

TO and JS: performed experiments, analyzed and evaluated data and drafted figures; PS performed and evaluated proteomics experiments and statistical analysis, SL: contributed the LC-MS metabolome data, MK: performed PHB quantification, BM: supervised proteome study; MH: supervised and evaluated metabolomics analysis and interpreted data; KF: designed and supervised the study, evaluated and interpreted data and wrote manuscript with the input from all authors.

