## Supplementary Material (Figures and Tables) for "The Novel PII-Interacting Regulator PirC (Sll0944) Identifies 3-Phosphoglycerate Mutase (PGAM) as Central Control Point of Carbon Storage Metabolism in Cyanobacteria"

### Table S1

#### Table S1 – Used Organisms and strains in this study

| Organism | Strain | Genotype | Purpose |
|---|---|---|---|
| *E. coli* | Top10 | F- mcrA Δ(mrr-hsdRMS-mcrBC) Φ80lacZΔM15 ΔlacX74 recA1 araD139 Δ( araleu)7697 galU galK rpsL (StrR) endA1 nupG | Molecular Cloning |
| *E. coli* | NEB10β | Δ(ara-leu) 7697 araD139 fhuA ΔlacX74 galK16 galE15 e14- Φ80dlacZΔM15 recA1 relA1 endA1 nupG rpsL (StrR) rph spoT1 Δ(mrr-hsdRMS-mcrBC) | Molecular Cloning |
| *E. coli* | Lemo21(DE3) | fhuA2 [lon] ompT gal (λ DE3) [dcm] ΔhsdS/ pLemo(CamR) λ DE3 = λ sBamHIo ΔEcoRI-B int::(lacI::PlacUV5::T7 gene1) i21 Δnin5<br>pLemo = pACYC184-PrhaBAD-lysY | Protein Expression |
| *E. coli* | Lemo21(DE3)<br>+ pJS15 | Lemo21(DE3)<br>+ pJS15 | Expression of strep-Tagged PII protein |
| *E. coli* | Lemo21(DE3)<br>+ pJS22 | Lemo21(DE3)<br>+ pJS22 | Expression of $His_8$-Tagged PirC |
| *E. coli* | Lemo21(DE3)<br>+ pJS26 | Lemo21(DE3)<br>+ pJS26 | Expression of $His_8$-Tagged PirC |
| *E. coli* | Lemo21(DE3)<br>+ pJS27 | Lemo21(DE3)<br>+ pJS27 | Expression of strep-Tagged PirC |
| *E. coli* | Lemo21(DE3)<br>+ pET-28a(+)-PGAM | Lemo21(DE3)<br>+ pET-28a(+)-PGAM | Expression of $His_6$-Tagged PGAM |
| *Synechocystis* sp. PCC 6803 | wild type glucose sensitive | WT | Background strain, control |
| *Synechocystis* sp. PCC 6803 | ΔPII | ssl0707::$Spec^R$ | Control strain |
| *Synechocystis* sp. PCC 6803 | Δ*pirC* | *pirC*::$Kan^R$ | Characterization of sll0944 KO mutants |
| *Synechocystis* sp. PCC 6803 | Δ*pirC*::*pirC* | *pirC*::$Kan^R$ pVZ322-pirC | Complementation of sll0944 knock out |
| *Synechocystis* sp. PCC 6803 | Δ*pirC*::*pirC*-mCitrine | Δ*pir*C pVZ322-pirC-mCitrine | Localization of PirC in Δ*pirC* and Co-immunoprecipitation |
| *Synechocystis* sp. PCC 6803 | Δ*pirC*::*pirC*-eGFP | Δ*pir*C pVZ322-*pirC*- eGFP | Localization of PirC in Δ*pirC* and Co-immunoprecipitation |

Table S2

Table S2 – Used Primer in this study

| #No | Primer | Sequence (5'-3') |
|---|---|---|
| #1 | pUC19-pirCUS_fw | GTTTTCCCAGTCAGACGTTGTAAAACGACGGCCAGTGAATTGTCGGCTGAAATTCATC |
| #2 | pirCUS-KanR_rev | GAATTGACATAAGCCTGTTCAATCAAATTTTTTGACTCTG |
| #3 | pirCUS-KanR_fw | CAGAGTCAAAAAATTTGATTCAACAGGCTTATGTCAATTC |
| #4 | KanR-pirCDS_rev | CCAATTCATTAATTCATTGGAGTTTGTAGAAACGCAAAAA |
| #5 | KanR-pirCDS_fw | GCCATCCTGACGGATGGCCTTTTTGCGTTTCTACAAACTCCAATGAATTAATGAATTGG |
| #6 | pirCDS-pUC19_rev | CGCCAAGCTTGCATGCCTGCAGGTCGACTCTAGAGGATCTGGAACATGGCTTCCCCTTTC |
| #7 | pET15b-pirC(correct)_fw | CATCATCATCACAGCAGCGGCCTGGTGCCGCGCGGCAGCATGTCGCAAATCTTGGACCC |
| #8 | pirC-pET15b_rev | CCTTTCGGGCTTTGTTAGCAGCCGGATCCTCGAGCATACTATGCGACAAGAGATTGAC |
| #9 | pASKIba5Plus-pirC_fw | CCACCCGCAGTTCGAAAAAGGCGCCGAGACCGCGGTCCCGATGTCGCAAATCTTGGACCC |
| #10 | pASKIba5Plus-pirC_fw | GGTCGACCTCGAGGGATCCCCGGGTACCGAGCTCGAATTCTATGCGACAAGAGATTGAC |
| #11 | Slr1945-NdeI-fw | CATATGATGGCAGAGGCACCGATCGCC |
| #12 | Slr1945-BamHI-rv | GGATCCCTAACGGGAGAGATTGACCGG |
| #13 | pVZmCit_pirC_fwd | CTGCAGGAGCAGAAGAGCATAC |
| #14 | pVZmCit_ pirC_rev | AGCCACTAAGGATTGGGAAG |
| #15 | pTO201_eGfp_fwd | ACACGAGTCCGAGGATATGACTTCCCAATCCTTAGTGGCTATGAGTAAAGGAGAAGAAC |
| #16 | pTO201_eGFP_rev | CTGGCTTTGCTTCCAGATGTATGCTCTTCTGCTCCTGCAGTTATTTGTATAGTTCATCCATGC |
| #17 | 6803 pirCKOcheck_fw | TGGCATGGCCTAAGTATTCC |
| #18 | 6803 pirCKOcheck_rev | GCGTTCTGCAGGGGATTACC |
| #19 | pVZ322seq_fw | CCTGGCTTTGCTTCCAGATG |
| #20 | pVZ322seq_rev | TGCCCGGATTACAGATCCTC |
| #21 | seq_pTO201_fwd | CAATGCTTTGCGAGATACCC |
| #22 | seq_pTO201_rev | AGCTCCATAGGCCGCTTTC |

### Table S3

Table S3 – Used Plasmids in this study

| Plasmid | Purpose | Source |
|---|---|---|
| pJS15 | Expression of Strep-Tagged PII protein (Ssl0707) in E. coli | Scholl et al (2020) |
| pJS22 | Expression of $His_8$-Tagged PirC (Sll0944) in E. coli T7-strains | This study |
| pJS26 | Expression of $His_8$-Tagged PII protein (Ssl0707) in E. coli T7-strains | Scholl et al (2020) |
| pJS27 | Expression of strep-Tagged PirC (Sll0944) in E. coli | This study |
| pJS31 | $Kan^R$ deletion of the pirC gene in Synechocystis sp. PCC 6803 | This study |
| pET-28a(+)-PGAM | Expression $His_8$-Tagged PGAM (Slr1945) in E. coli T7-strains | This study |
| pVZ322-pirC-comp | Complementation of pirC deletion | Klotz (2017) |
| pVZ322-mCitrine-pirC-comp | Complementation of pirC deletion tagged to mCitrine protein | Klotz (2017) |
| pTO201 | Complementation of pirC deletion tagged to eGFP protein | This study |

### Figure S1

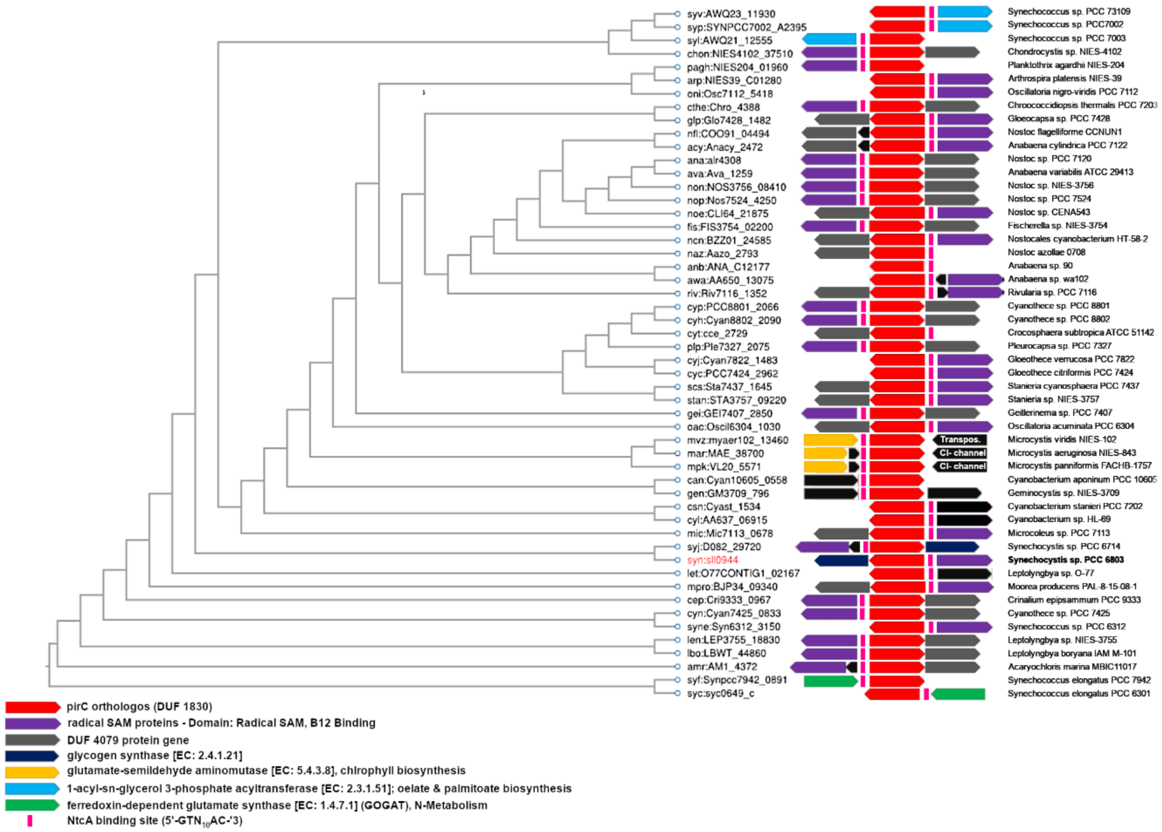


**Fig. S1 - Cluster comparison of 53 different Cyanobacterial PirC orthologous.** Detected by a Smith-Waterman (SW) algorithm based SSEARCH for bidirectional best hits of the KEGG SSDB (Sequence Similarity Database) with a SW-Score threshold of 100. Red arrows represents the pirC orthologous with direction. In front of each pirC gene a predicted NtcA binding site is present (5'-GTN10AC-'3, pink bars). In 67 % of the cases the pirC are next to genes encoding radical SAM-like proteins (purple arrows).

Figure S2

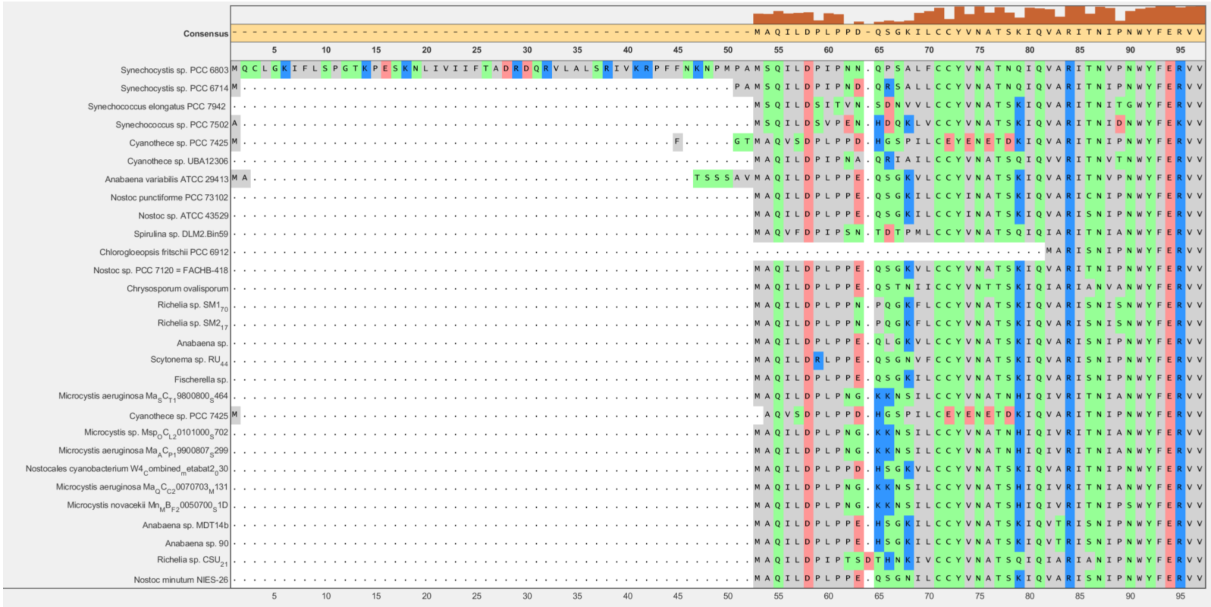


**Fig. S2 – Multiple alignment of PirC orthologs of different cyanobacteria.** It is shown that the first 52 AA are only present in *Synechocystis* sp. PCC 6803 PirC (first row). The picture shows extract of alignment of 29 different cyanobacterial orthologs of a multiple alignment of 74 different orthologs.

Figure S3

A

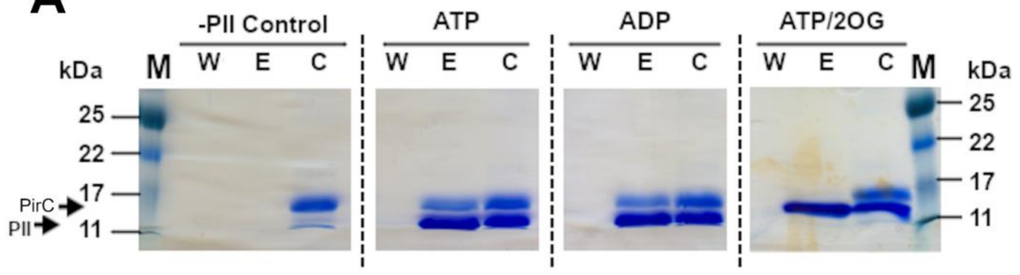


B

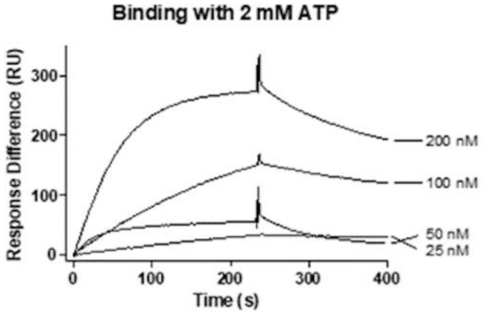


**Figure S3 - SDS-PAGE of in-batch pulldown assays in presence of 10 mM $MgCl_2$ and either 2 mM ATP, 2 mM ADP, or 2 mM ATP/2-OG.** The first column represents the negative control without $P_{II}$**.** PirC was co-eluted with $P_{II}$ attached to strep-tactin XT coated magnetic beads in presence of ATP and ADP, observable by to bands. However, there is no band in presence of ATP/2-OG. In the controls without $P_{II}$, no elution of PirC was observed. 1: Marker; W: washing fractions; E: elution fractions; C: control (reaction mix before the pulldown was performed).

Figure S4

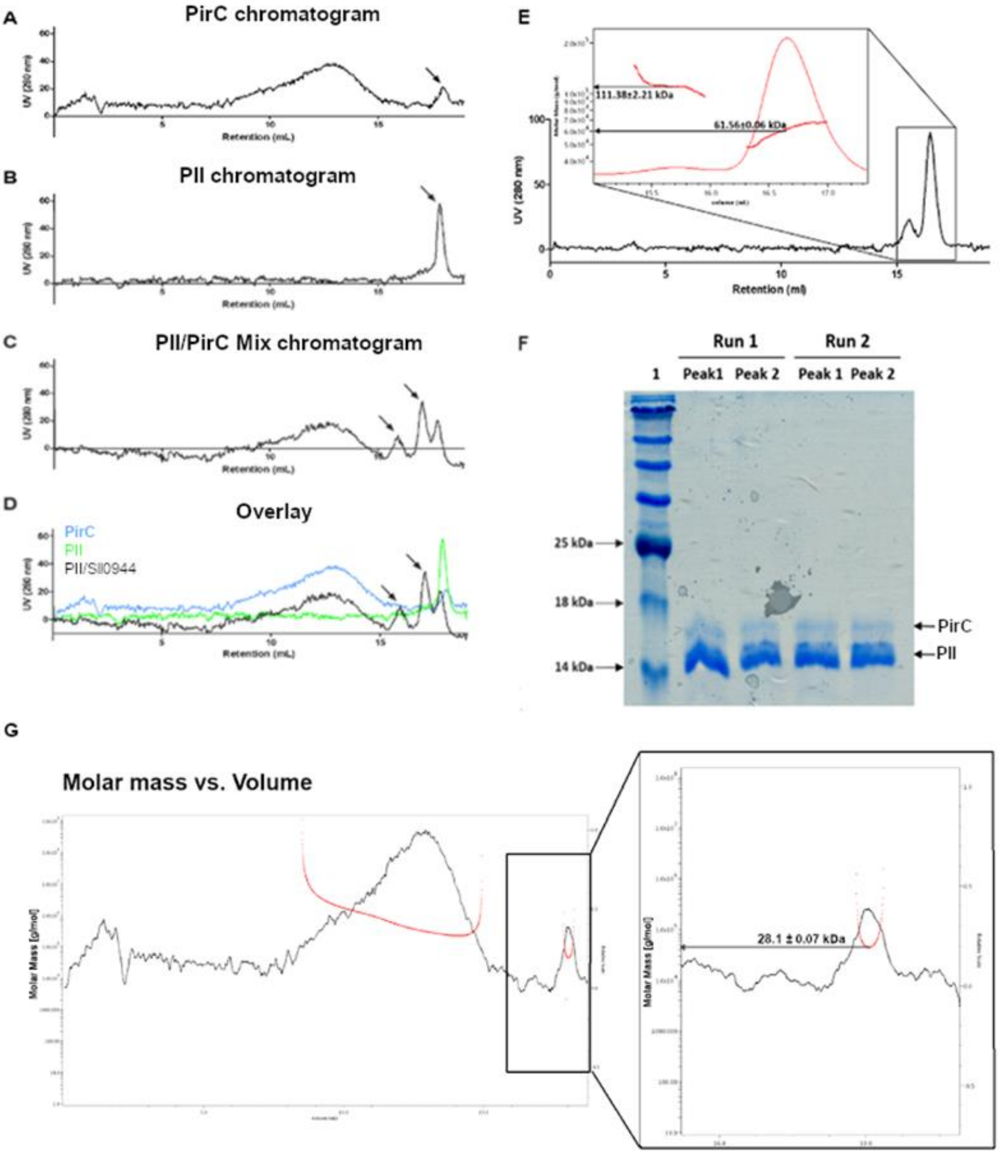


**Fig. S4 - SEC-MALS elution profiles of recombinant *Synechocystis* $H_6$-PirC and Strep-PII. (A)** The recombinant PirC elutes in high oligomeric states with a distinct fraction being dimeric (apparent molecular weight 28.10±0.07 kDa; see arrow); **(B)** PII elutes in a single peak as a trimer (45.7±0.09 kDa; see arrow); **(C)** In the combined run with both, PirC and PII, in a 1:1 mixture (20 nmol of each protein in a volume of 100 µL), in addition to the PirC high oligomeric peak and the PII peak, to new peaks appear in the elution profile (see arrows), with the larger peak yielding an apparent molecular weight of 58.5±0.22 kDa and the smaller peak yielding an apparent weight of 113.41±2.46 kDa. This corresponds to a complex made of one PII trimer and one PirC monomer, and a dimer of this complex, respectively. **(D)** Overlay of all three elution profiles with the PII/PirC complex peaks (marked with arrows). **(E).** Two peaks appear in the elution profile, with the larger peak yielding an apparent molecular weight of 61.56±0.06 kDa and the smaller peak yielding an apparent weight of 111.38±2.21 kDa. This corresponds to a complex made of one PII trimer and one PirC monomer, and a dimer of this complex, respectively. Representative graph of three runs; F**.** SDS-PAGE analysis of both peak fractions from both runs. Both peaks contained PirC and PII in both runs. **(G)** SEC-MALS elution profiles of recombinant *Synechocystis* $H_6$-PirC.

### Figure S5

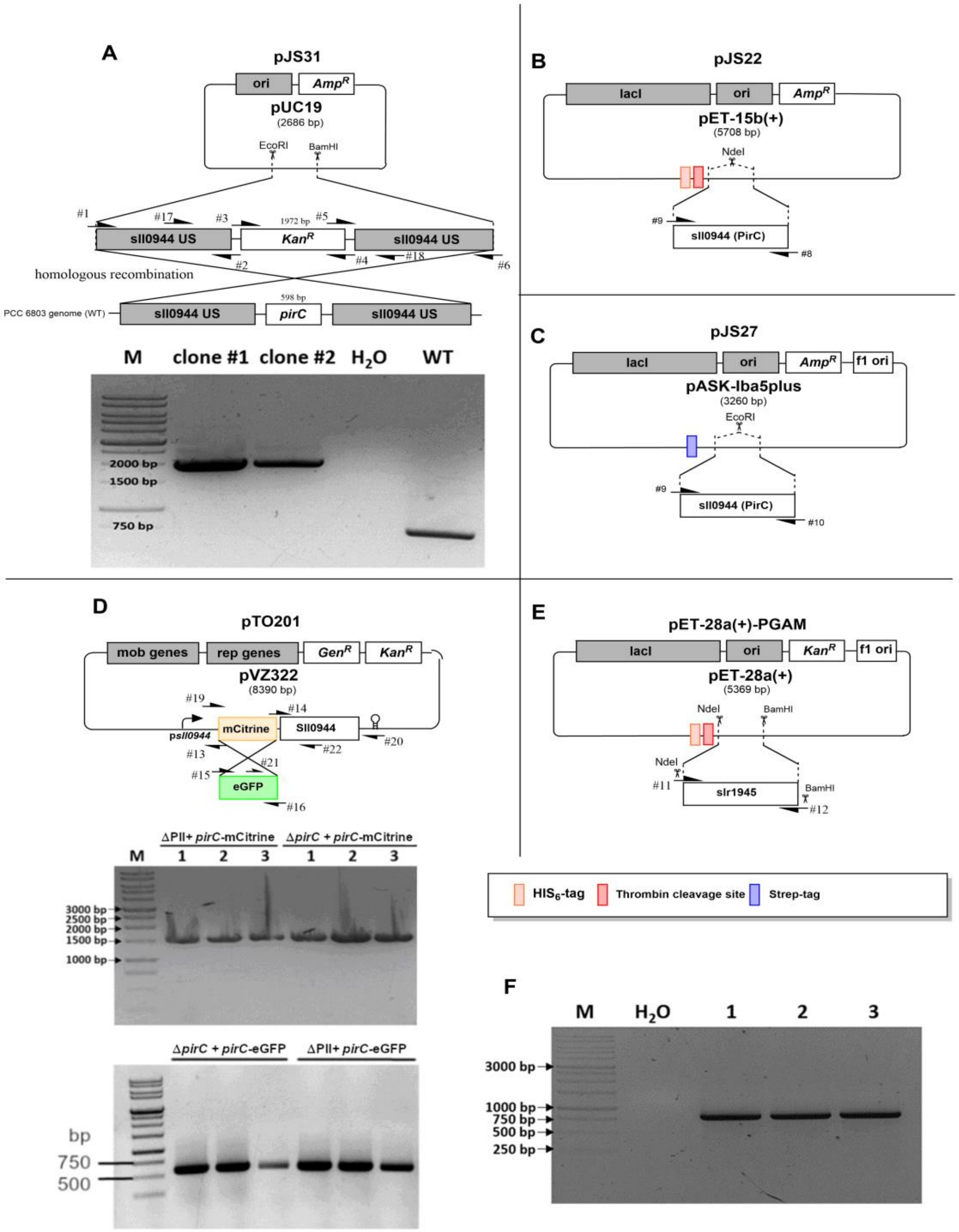


**Fig. S5 - Plasmid Construction and mutant check in *Synechocystis* sp. PCC 6803. (A)** Plasmid construction, gene deletion technique, manufactured by Gibson Assembly and colony PCR check of *Synechocystis* Δ*pirC* gene using primer pair #17/18 **(B)** Plasmid construction pJS22 for His$_6$-PirC expression in *E. coli* , manufactured by Gibson Assembly **(C)** Plasmid construction pJS27 for Strep-PirC expression in *E. coli* , manufactured by Gibson Assembly **(D)** Plasmid construction pTO201 for eGFP-PirC expression in *Synechocystis*, manufactured by Gibson Assembly as well as colony PCR of successful transformation of pVZ322-mCitrine-*pirC*-comp and pTO201.**(E)** Plasmid construction pET-28a(+)-PGAM for His6-PirC expression in *E. coli.* Manufactured by Restriction/ligation method **(F)** colony PCR of successful transformation of pVZ322-pirC-comp

Figure S6

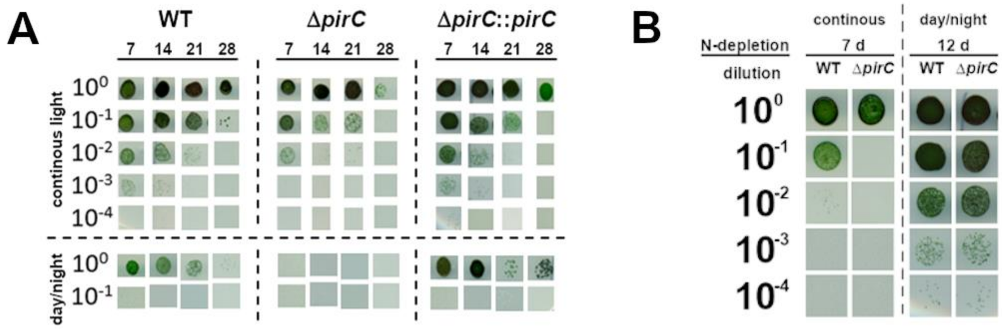


**Figure S6 - Viability drop agar test of *Synechocystis* WT, Δ*pirC* and Δ*pirC::pirC*.** **(A)** Drops were placed on the plates after 7, 14, 21, and 28 days of chlorosis. For each day, two plates were inoculated, and one was grown under continuous light while the other was grown in the day/night cabinet. Scans of the plates were made after 10 days (continuous light) or after 14 days (day/night) of growth. Each column represents a biological replicate. **(B)** Drops were placed on the plates after 7 or 12 days of chlorosis either under continuous or day/night illumination. Scans of the plates were made after 7 days. Each column represents a biological replicate.

Figure S7

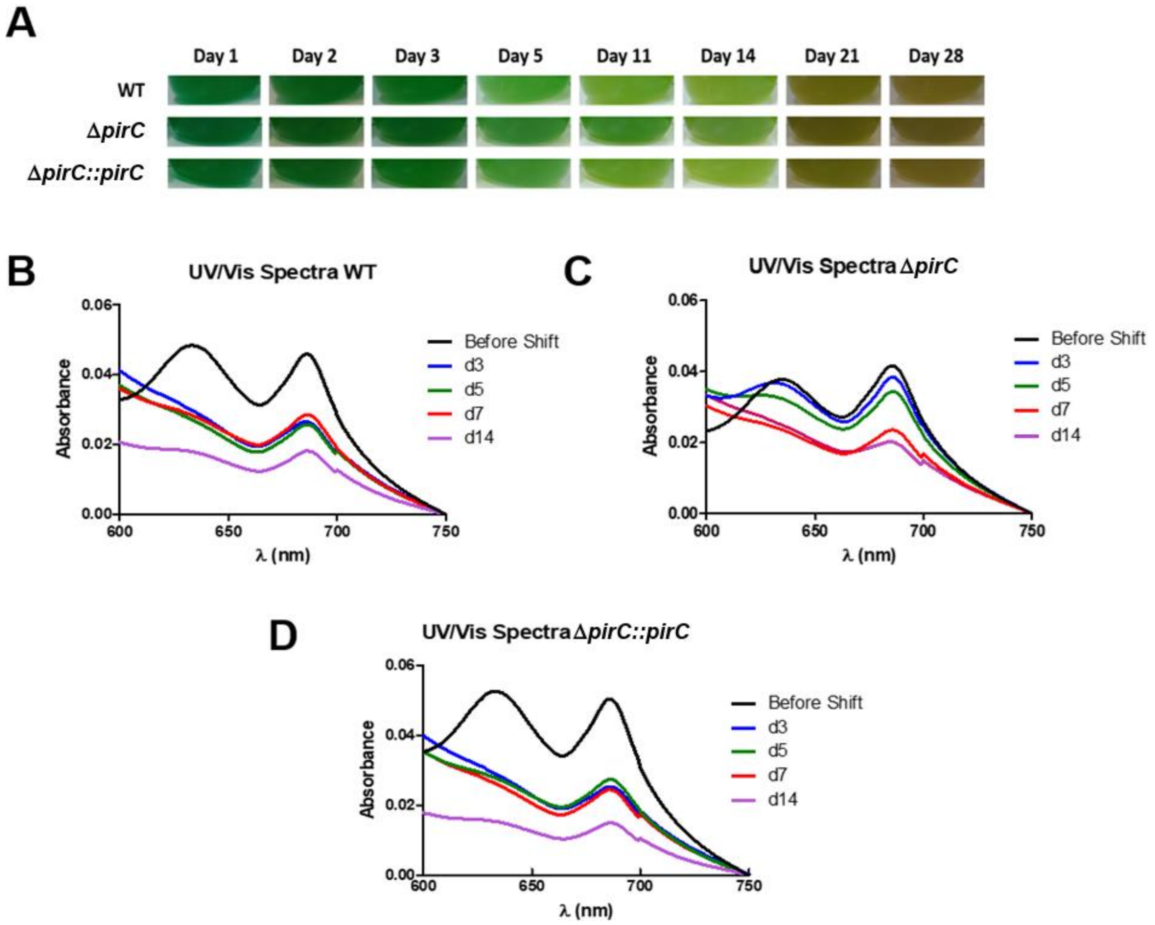


**Fig. S7 - Chlorosis due to nitrogen depletion of *Synechocystis* wild type, Δ*pirC*, and Δ*pirC::pirC* in the day/night cabinet. (A)** Photographs showing depigmentation of representative cultures from the three replicates; **(B)** UV/Vis spectra (mean of three replicates) of the wild type; **(C)** UV/Vis spectra (mean of three replicates) of the Δ*pirC* mutant; **(D)** UV/Vis spectra (mean of three replicates) of the complemented mutant Δ*pirC::pirC*.

Figure S8

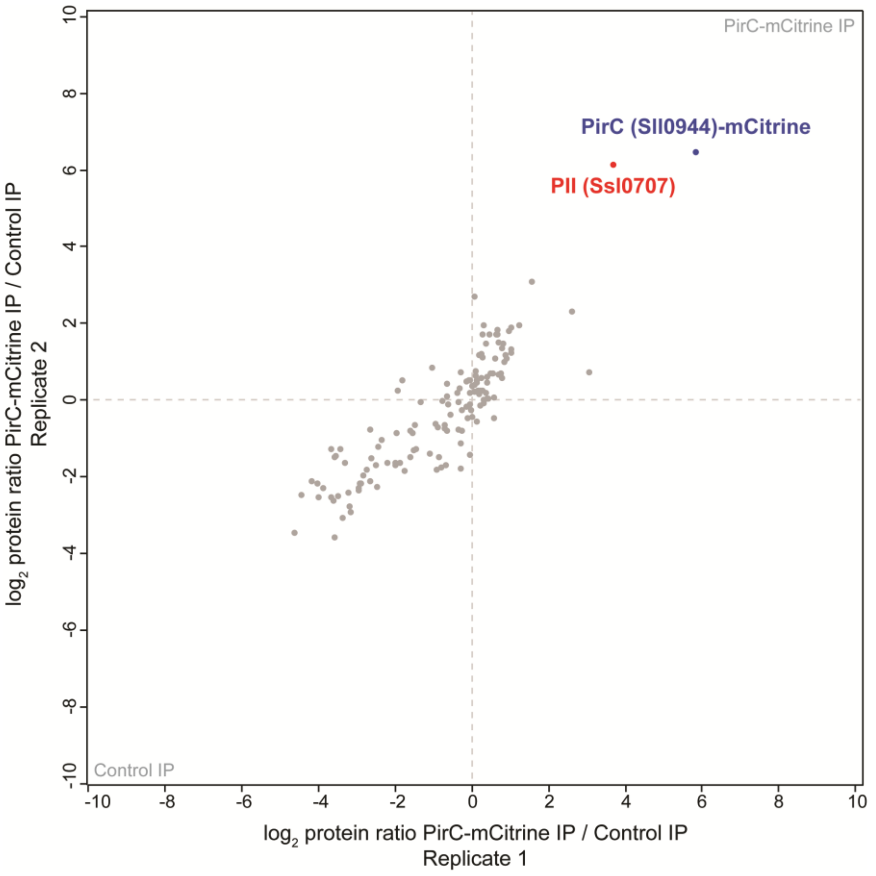


**Fig. S8 - Validation of the interaction of the PirC-mCitrine construct with PII.** The newly designed PirC-mCitrine fusion protein was tested in anti-GFP immunoprecipitation (IP) experiments for interaction with the PII protein at conditions of low ATP and 2-OG levels, as revealed in reverse IP experiments against a tagged PII construct (Watzer at al. 2019). The scatterplot of two independent PirC-mCitrine IP experiments confirms a significant co-enrichment (p-value=0.01) of PirC and PII, indicated in blue and red, respectively. The IP was performed with crude cell extracts from nitrogen-starved cells of the Δ*pirC*::*pirC-mCitrine* strain, without addition of key metabolites. Extracts were either incubated with GFP-Trap coated magnetic agarose beads (PirC-mCitrine IP) or protein A/G agarose beads coated with an unrelated antibody (Control IP). IP eluates were differentially labeled by dimethlytion labeling and analyzed by high accuracy LC-MS/MS. MS data was processed and analyzed as described elsewhere (Boersema et al 2009, Watzer et al. 2019).

Watzer, B., P. Spat, N. Neumann, M. Koch, R. Sobotka, B. Macek, O. Hennrich and K. Forchhammer (2019). "The Signal Transduction Protein P-II Controls Ammonium, Nitrate and Urea Uptake in Cyanobacteria." Frontiers in Microbiology 10.

Boersema, P. J., R. Raijmakers, S. Lemeer, S. Mohammed and A. J. R. Heck (2009). "Multiplex peptide stable isotope dimethyl labeling for quantitative proteomics." Nat. Protocols 4(4): 484-494.

Figure S9

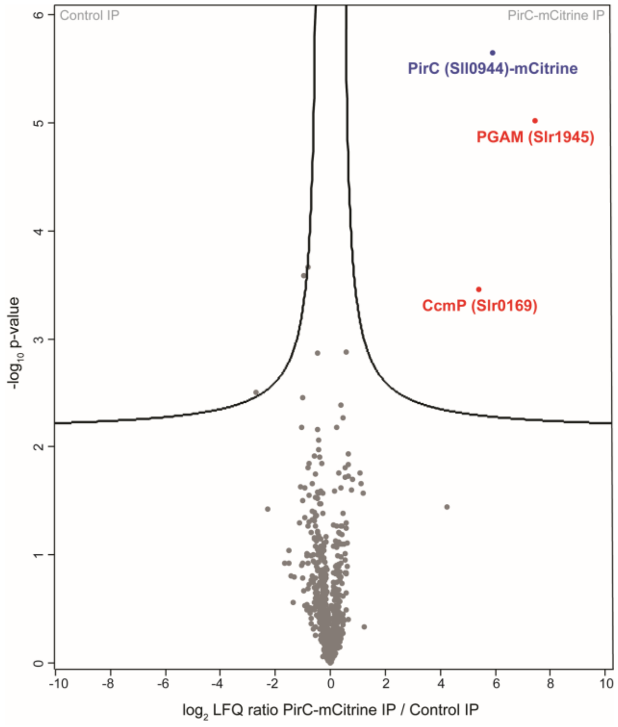


**Fig. S9 - The PirC interactome at nitrogen starvation conditions.** Volcanoplot of three independent PirC-mCitrine immunoprecipitation (IP) experiments displays co-enriched proteins. Crude cell extracts from the nitrogen-starved Δ*pirC*::*pirC-mCitrine* strain were incubated with either GFP-Trap coated magnetic agarose beads (PirC-mCitrine IP) or non-coated beads (Control IP) in presence of ATP, 2-OG and $Mg^{2+}$(each 2 mM). IP eluates were analyzed by high accuracy LC-MS/MS using label-free quantification to calculate protein enrichment ratios. Significantly enriched proteins were defined by *t*-test (FDR=0.01; S0=0.1) and are indicated in blue (PirC-mCitrine) or red (co-enriched proteins).

Figure S10

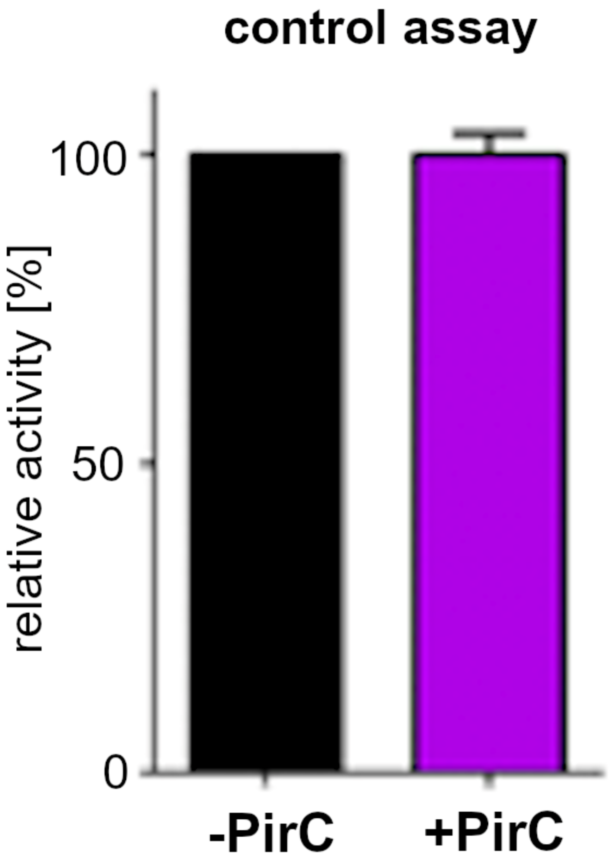


**Fig. S10 – Control assay of coupled enzymes.** Black bar represents the mean of triplicates with SD of control assay without PirC addition. Purple bar represents the mean of triplicates with SD of control assay with the addition of 600 nM PirC. In the assay 0.625 mM 2-PGA was added to the assay.

#### Figure S11

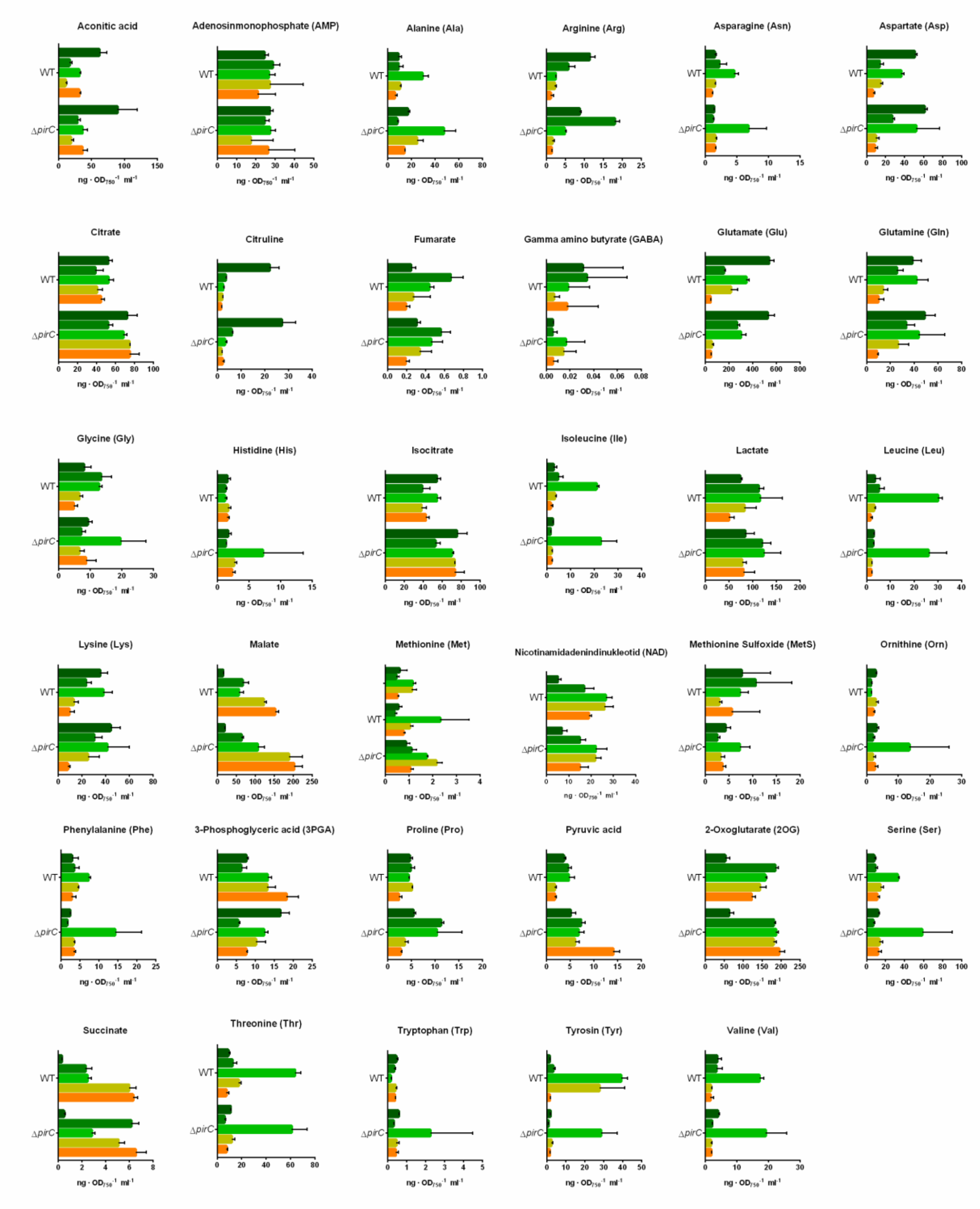


**Fig. S11 – LC-MS analysis of the hole spectrum of measured compounds.** On x-axis the concentration of the compounds is shown in ng · $OD_{750}^{-1}$ · $ml^{-1}$. Each bar represents the mean of two technical replicates of two biological replicates. The Error bar represents the SD of the measurement.

Figure S12

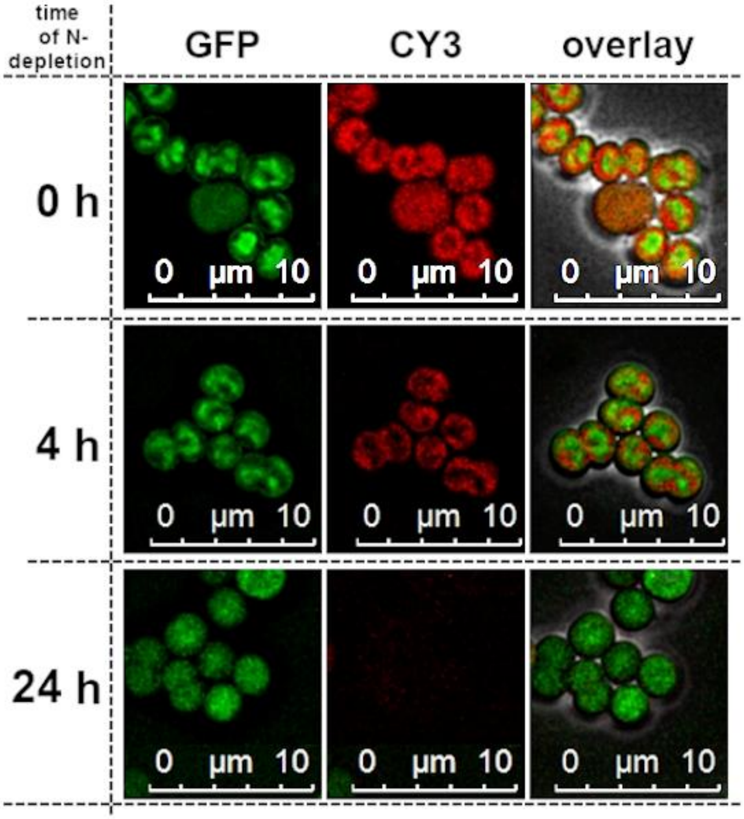


**Figure S12 - Intracellular localization of the PirC-eGFP fusion protein in *Synechocystis* sp. PCC 6803.** Change of the intracellular localization of the PirC-eGFP signal in *Synechocystis* after nitrogen depletion. Representative pictures of three biological replicates. Directly after the shift to nitrogen depleted conditions, the signal is localized centrally in the cytoplasm (0h) with no clear change after 4 h (4 h). However, the signal is more distributed throughout the cell after 24 h.
